# Global analysis of the zinc homeostasis network in *Pseudomonas aeruginosa* and its gene expression dynamics

**DOI:** 10.1101/2021.05.28.446167

**Authors:** Verena Ducret, Melina Abdou, Catarina Goncalves Milho, Sara Leoni, Oriane Martin--Pelaud, Antoine Sandoz, Inés Segovia Campos, Mary-Lou Tercier-Waeber, Martina Valentini, Karl Perron

## Abstract

Zinc is one of the most important trace elements for life and its deficiency, like its excess, can be fatal. In the bacterial opportunistic pathogen *Pseudomonas aeruginosa*, Zn homeostasis is not only required for survival, but also for virulence and antibiotic resistance. Thus, the bacterium possesses multiple Zn import/export/storage systems. In this work, we determine the expression dynamics of the entire *P. aeruginosa* Zn homeostasis network at both transcript and protein levels. Precisely, we followed the switch from a Zn-deficient environment, mimicking the initial immune strategy to counteract bacterial infections, to a Zn-rich environment, representing the phagocyte metal boost used to eliminate an engulfed pathogen. Thanks to the use of the NanoString technology, we timed the global silencing of Zn import systems and the orchestrated induction of Zn export systems. We show that the induction of Zn export systems is hierarchically organized as a function of their impact on Zn homeostasis. Moreover, we identify PA2807 as a novel Zn resistance component in *P. aeruginosa* and highlight new regulatory links among Zn-homeostasis systems. Altogether, this work unveils a sophisticated and adaptive homeostasis network, which complexity is key in determining a pathogen spread in the environment and during host-colonization.

## Introduction

Zinc (Zn), commonly found as the divalent cation Zn^2+^, is a trace element primarily involved as a cofactor of many enzymes and therefore essential for life. It is considered as the second most important trace metal after iron (Fe) (1). In prokaryotes, Zn is principally found bound to proteins (approximately 6% of the proteome) and the free cellular fraction is kept at a concentration as low as femtomolar (2). In excess, this metal becomes toxic, especially by competing with other trace elements, giving rise to protein mismetallization (3). The equilibrium of cellular Zn concentration is therefore tightly regulated and numerous systems involved in Zn homeostasis (and that of other divalent cations) are well-described in several bacteria (4).

The toxic properties of Zn are widely exploited in the context of infection control. Upon bacterial invasion, one of the host defense mechanisms is the sequestration of essential metal ions, in particular Fe, Zn and manganese (Mn), by proteins and molecules, leading to so called “nutritional immunity” (5-7). Conversely, during phagocytosis, toxic concentrations of Zn and copper (Cu) are delivered into the phagolysosome, participating in the destruction of the pathogen (8). Thus, to successfully infect, microorganisms not only must adapt to environments of Zn deprivation or excess, but also rapidly alternate between these two extreme conditions. The opportunistic Gram-negative pathogen *Pseudomonas aeruginosa* is an ubiquitous and versatile bacterium that possesses a huge armada of systems for Zn homeostasis (Fig 1A). *P. aeruginosa* is responsible for a wide array of severe infections, particularly in cystic fibrosis and immunocompromised patients (9), and Zn homeostasis systems are highly relevant to its pathogenicity (10). Notably, they modulate virulence by acting on quorum sensing (11) as well as antibiotic resistance by repressing the expression of the route of entry for carbapenem antibiotics (12).

**Figure 1:**
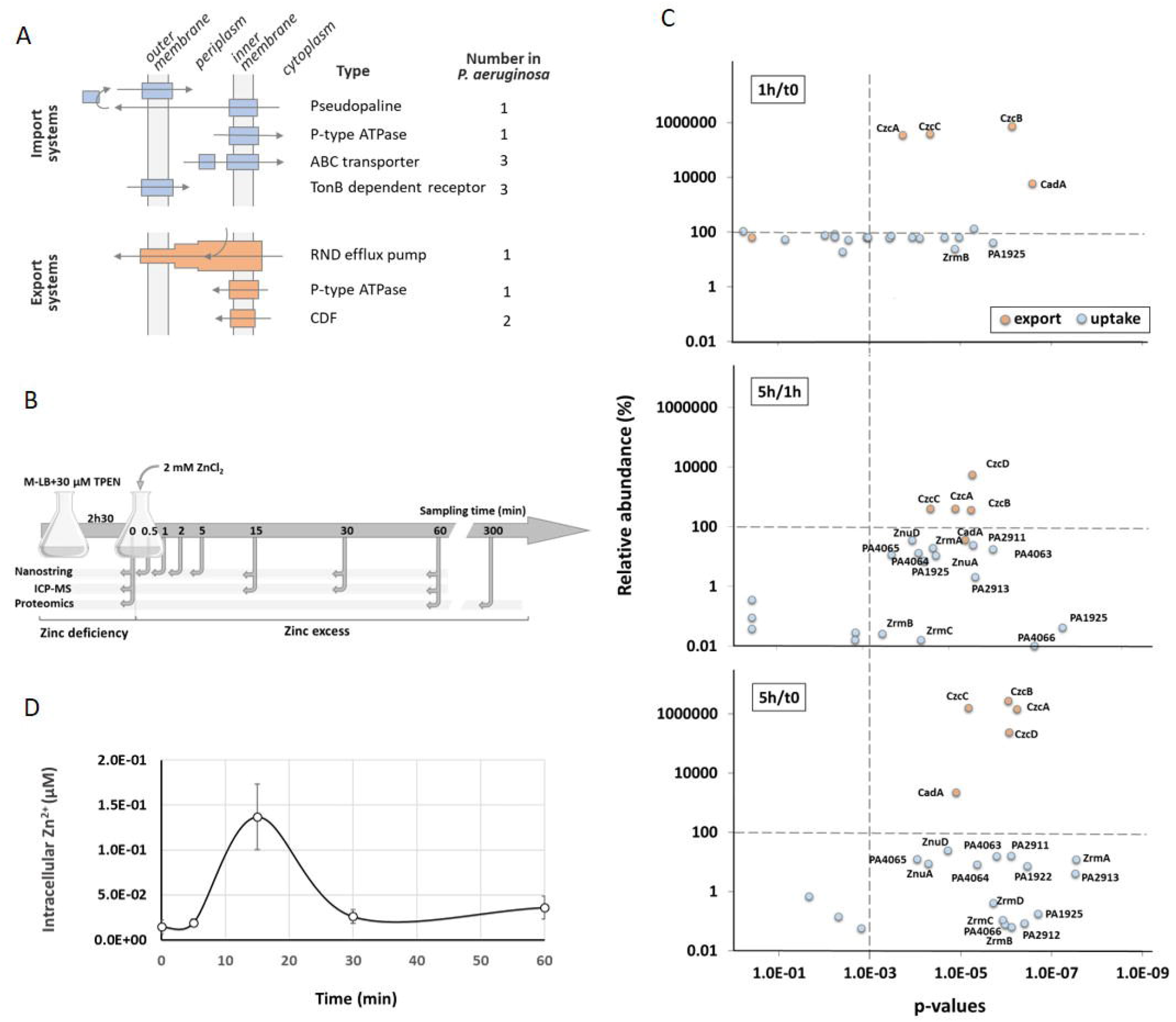
Dynamics of Zn homeostasis in *P. aeruginosa*. (A) The four families of Zn import systems (in blue) and the 3 families of Zn export systems (in orange) are represented. Their cellular localization and the number of each representative are indicated. (B) Schematic representation of the experimental setup and the times of sampling. (C) Relative abundance of the zinc transport systems. Graphs represent the correlation plots between the percentage of peptides detected (y axis) after 1h compared to t0 (upper panel), 5h compared to 1h (middle panel) or 5h compared to t0 (lower panel) of Zn treatment, and their corresponding statistical values (x axis). The p-values lower than 10^−3^ (to the right of the vertical dotted line) were considered as statistically significant. The average values obtained for t0 (or 1h for the middle panel) correspond to 100%. (D) Changes in intracellular zinc concentration after addition of 2 mM ZnCl_2_ to the medium. The metal concentrations in μM were determined over time in the wild type strain by ICP-MS.

Numerous systems are involved in the import of Zn into *P. aeruginosa* (13). At first, the TonB-dependent receptors ZnuD and PA2911 interact with its energizing TonB-ExbBD protein complex to translocate extracellular Zn into the periplasmic space (10). Subsequently, the metal crosses the cytoplasmic membrane via either the P-type ATPase HmtA or via an ATP-binding cassette (ABC) transporter such as ZnuABC, the most common Zn uptake pathway in bacteria (13). Revealed for the first time in *Escherichia coli*, this transporter consists of the solute-binding protein (SBP) ZnuA, which scavenges Zn in the periplasm with a high affinity and brings it to the inner membrane permease ZnuB. The energy required for this transport is provided by the cytoplasmic ATPase ZnuC (14).

Two additional ABC transporters have been described in *P. aeruginosa*: PA4063-PA4066 and PA2912-PA2914 (13). Although these transporters have been shown to be overexpressed under conditions of Zn limitation, they remain poorly characterized and their exact contribution to Zn uptake remains to be elucidated.

The ZrmABCD system has recently been designated as a new Zn acquisition system, involving a nicotianamine-related zincophore called pseudopaline by similarity to the *Staphylococcus aureus* staphylopine (15,16). The molecule is synthetized by the ZrmB and ZrmC cytoplasmic enzymes and then released outside the cell by the EamA-like transporter ZrmD. Internalization of the Zn-pseudopaline complex is mediated by the TonB-dependent receptor ZrmA, in a siderophore-like manner (17). Though many gaps remain to be filled at the mechanical level, the importance of this system under conditions of Zn limitation and its involvement in bacterial pathogenicity appear evident, especially in a cystic fibrosis context (18).

A considerable reservoir of Zn is represented by ribosomal proteins. Indeed, during nutrient deprivation, *P. aeruginosa* copes with metal deficit by expressing additional ribosomal paralogous proteins, annotated as C-, which have lost their Zn finger domain (10,19). A release of Zn probably results from this exchange phenomenon, that can be used by other proteins such as DNA polymerase, primase, etc. The *P. aeruginosa* PA3600-PA3601 operon encodes two C-paralogs, RpmE2 and RpmJ2, able to substitute in place and in function the ribosomal proteins RpmE and RpmJ respectively. Recent data from *E. coli* have shown that RpmE2 and RpmJ2 are of comparable efficiency to their Zn-ribbon paralogs (20). Such a strategy is also observed with the global transcription regulator DksA, involved in the stringent response. Under Zn-scarce conditions, the alternative protein DksA2, devoid of a Zn finger domain, is induced and takes over the functional relay from its paralog DksA (21,22).

Zn uptake and storage systems are regulated by the one-component regulator Zn uptake regulator (Zur) protein, that belongs to the ferric uptake regulatory (FUR) protein family. Zur senses the cytoplasmic concentration of the metal and its regulatory function is directly related to its ability to reversibly bind Zn (10,23). Under conditions of Zn sufficiency or excess, the regulator is found in a dimeric form containing four Zn atoms (Zur2-Zn4 conformation), which promotes DNA binding and ensures the repression of target genes by preventing RNA polymerase from initiating transcription. In *P. aeruginosa*, nine Zur binding sites were predicted, characterized by a 17-nucleotide palindromic motif overlapping the -10 of the promoter region (13).

*P. aeruginosa* is also very well-armed to counter high toxic concentrations of Zn. Three families of export systems have been described in this bacterium, including the CzcCBA efflux pump, a homolog of the system found in the metal-resistant bacterium *Cupriavidus metallidurans* (24). CzcCBA belongs to the RND (Resistance-Nodulation-Division) group of the HME (Heavy Metal Efflux) family, capable of expelling excess Zn, cadmium (Cd) and cobalt (Co) from the cytoplasmic or periplasmic compartments directly outside the cell (25). This pump is regulated by the CzcRS two-component system (TCS), where CzcS is the transmembrane sensor and CzcR is the response regulator. CzcS detects and becomes active under conditions of excess periplasmic Zn or Cd. It then activates CzcR by phosphorylation, which in turn acts as a transcriptional activator of the efflux system, ensuring detoxification of the cell. Additionally, CzcRS TCS has been shown to activate its own transcription, promoting a positive regulation loop, but also to directly repress the OprD porin, thus rendering the bacterium resistant to carbapenem antibiotics (11,12).

Moreover, the bacterium possesses two cation diffusion facilitators (CDF), CzcD and YiiP. These systems act as a homodimer or heterodimer to expel Zn from the cytoplasm to the periplasm by way of a proton gradient. They also appear to be involved in maintaining membrane integrity, which would explain why mutants deleted for these CDFs are more sensitive to several antibiotics (26).

A final layer consists of the P-type ATPase CadA, which is also responsible for detoxifying the cytoplasm by expelling Zn to the periplasm. In *P. aeruginosa*, CadA was first described as a system involved in Cd resistance (27). Similar to the *E. coli* ZntA, this protein has recently been shown to be essential for Zn resistance (28). CadA is regulated by CadR that belongs to the MerR family of response regulators. CadR is constitutively expressed and permanently located on the *cadA* promoter. The activity of this regulator is directly linked to its ability to bind Zn since it plays the dual role of repressor and activator of *cadA* transcription, depending on whether it is in conditions of Zn limitation or excess respectively (28).

Numerous strategies allowing Zn homeostasis have been described and characterized in *P. aeruginosa*. All these systems serve to ensure the survival of the bacterium in environments that are limited or contaminated with Zn and, interestingly, make the link between metallostasis, virulence and antibiotic resistance. Knowledge of how these systems interact with each other, however, is scarce.

Our lab previously highlighted a dynamic in the expression of Zn export systems in *P. aeruginosa*. Indeed, we had shown that the P-type ATPase CadA was not only the first bulwark against a boost in Zn, but also facilitated the induction of the major export system, the CzcCBA efflux pump (28). These results support the hypothesis that the different systems involved in Zn resistance are not redundant, but follow a precise strategic plan, guaranteeing a rapid adaptation of the cell to variations in Zn concentrations.

The purpose of this study was to follow up on the “strategic plan” hypothesis and assess the dynamics of all the systems involved in Zn homeostasis (import/export/storage) when the bacterium transitions from starvation to surfeit, as it happens during an infection. Using the NanoString technology (29), we could quantify transcripts of all Zn homeostasis systems, rather than following a single element of the system, and follow their dynamics over time. While we clearly observed a global import systems repression, a hierarchical induction of the export systems appeared. An additional novel partner of the Zn resistance, PA2807, a CzcE-like protein, was also discovered and characterized. Finally, the rapid induction of carbapenem resistance via OprD porin repression in the presence of Zn was precisely analyzed at the transcriptomic and proteomic levels.

## Results

In order to monitor the dynamics of Zn transport systems when *P. aeruginosa* switches from a situation of Zn starvation to Zn excess, we designed an experimental procedure described in Fig 1B. Briefly, *P. aeruginosa* PAO1 strain was cultivated in a Zn-depleted medium (M-LB) containing 30 µM TPEN (*N,N,N*′,*N*′-tetrakis(2-pyridinylmethyl)-1,2-ethanediamine) to chelate all residual Zn (23). Then, after 2h30 of growth at 37°C, 2 mM ZnCl_2_ (final concentration) was added to the culture. Previous Zn addition (t0) and at different times afterwards, cells were sampled for proteomic, transcriptomic and Zn content analysis.

At protein levels we observed, as expected, a significant increase (6.1×10^3^ to 7.3×10^5^ %) of all export systems during the first hour of high Zn exposure, while surprisingly the level of only two proteins involved in Zn import, ZrmB and PA1925, decreased more than 50% (Fig 1C, Table S1). Indeed, the large majority of uptake systems significantly dropped only between 1 to 5h after Zn addition. During this time frame, the amount of the CadA P-type ATPase strongly decreased as well, in agreement with our previous observations on its role of first-aid sentinel in case of a sudden Zn excess (28). The 5h-period was also marked by the increasing of the CzcD protein. This very late induction took place after the CzcCBA efflux rise and may explain why this CDF was not essential for Zn resistance(28).

When looking at the intracellular Zn concentration over time, the values changed from 0.014 to 0.14 µM in 15 min, then returned to equilibrium at a concentration of approximately 0.04 µM after 60 min (Fig 1D). It thus appears that the rapid induction of efflux systems prevails over the slow repression of import systems and it is sufficient to counteract the Zn overflow.

Finally, to decipher early events of regulation of the Zn homeostasis network, we directly monitored the transcript levels of genes involved in Zn transport using the NanoString technology (29). This method allows the direct counting of several mRNAs simultaneously using digital bar-coded probe pairs. Twenty-five oligonucleotide probes targeting mRNAs of the known Zn import/export/storage systems of *P. aeruginosa* were designed (Table 1). In addition, four probes targeting housekeeping genes were also designed for data normalization (Table S2). In case of a system encoded by operon, we selected at least one gene probe to represent the whole transcription unit (e.g. *czcA* and *czcC* were chosen to monitor the expression of the *czcCBA* operon). For this analysis, RNA was extracted at t0 and 30 seconds, 1, 2, 5, 15, 30 and 60 min (t0.5, t1, 2, 5, 13 and 60, respectively) after Zn addition (Fig 1B). The mRNA copy number for each of the target genes was then calculated in each sample, as shown in Table S3.

**Table 1:** List of the twenty-five genes monitored for zinc homeostasis analysis

| Pathway | Gene name | Gene ID | Product description |
| --- | --- | --- | --- |
| **Export** | *czcA* | PA2520 | Cation efflux transporter |
|  | *czcC* | PA2522 | Outer membrane protein |
|  | *cadA* | PA3690 | P-type ATPase transporter |
|  | *czcD* | PA0397 | Cation diffusion facilitator transporter |
|  | *yiiP* | PA3963 | Cation diffusion facilitator transporter |
| **Uptake** | *hmtA* | PA2435 | P-type ATPase transporter |
|  | *PA1922* | PA1922 | TonB-dependent receptor |
|  | *PA2911* | PA2911 | TonB-dependent receptor |
|  | *PA2914* | PA2914 | Permease of ABC transporter |
|  | *PA4063* | PA4063 | Solute-binding protein |
|  | *PA4065* | PA4065 | Permease of ABC transporter |
|  | *PA4066* | PA4066 | Solute-binding protein |
|  | *znuA* | PA5498 | Solute-binding protein |
|  | *znuB* | PA5501 | Permease of ABC transporter |
|  | *znuD* | PA0781 | TonB-dependent receptor |
|  | *zrmA* | PA4837 | TonB-dependent receptor |
|  | *zrmD* | PA4834 | Nicotianamine synthase |
| **Storage** | *dksA* | PA4723 | Supressor protein |
|  | *dksA2* | PA5536 | Supressor protein |
|  | *rpmE* | PA5049 | Ribosomal protein L31 |
|  | *rpmE2* | PA3601 | Ribosomal protein L31 |
|  | *rpmJ* | PA4242 | Ribosomal protein L36 |
|  | *rpmJ2* | PA3600 | Ribosomal protein L36 |
| **Others** | *PA2807* | PA2807 | Copper binding protein |
|  | *oprD* | PA0958 | Outer membrane porin |

## A) Zn export systems

### Dynamics of cadA and czcCBA expression

We could recently show that the two most important systems involved in Zn resistance in *P. aeruginosa* are the CadA P-Type ATPase and the CzcCBA RND efflux pump (28). These two systems interact together to produce an optimal response to Zn excess, *cadA* being rapidly induced while *czcCBA* expression being expressed successively, following a decrease in *cadA* expression (28). The *cadA* and *czcCBA* expression dynamic is also clearly visible in the NanoString experiment presented here (Fig 2A), thus confirming that this method is suitable for the fine monitoring of gene expression during the low-to-high Zn concentration transition.

**Figure 2:**
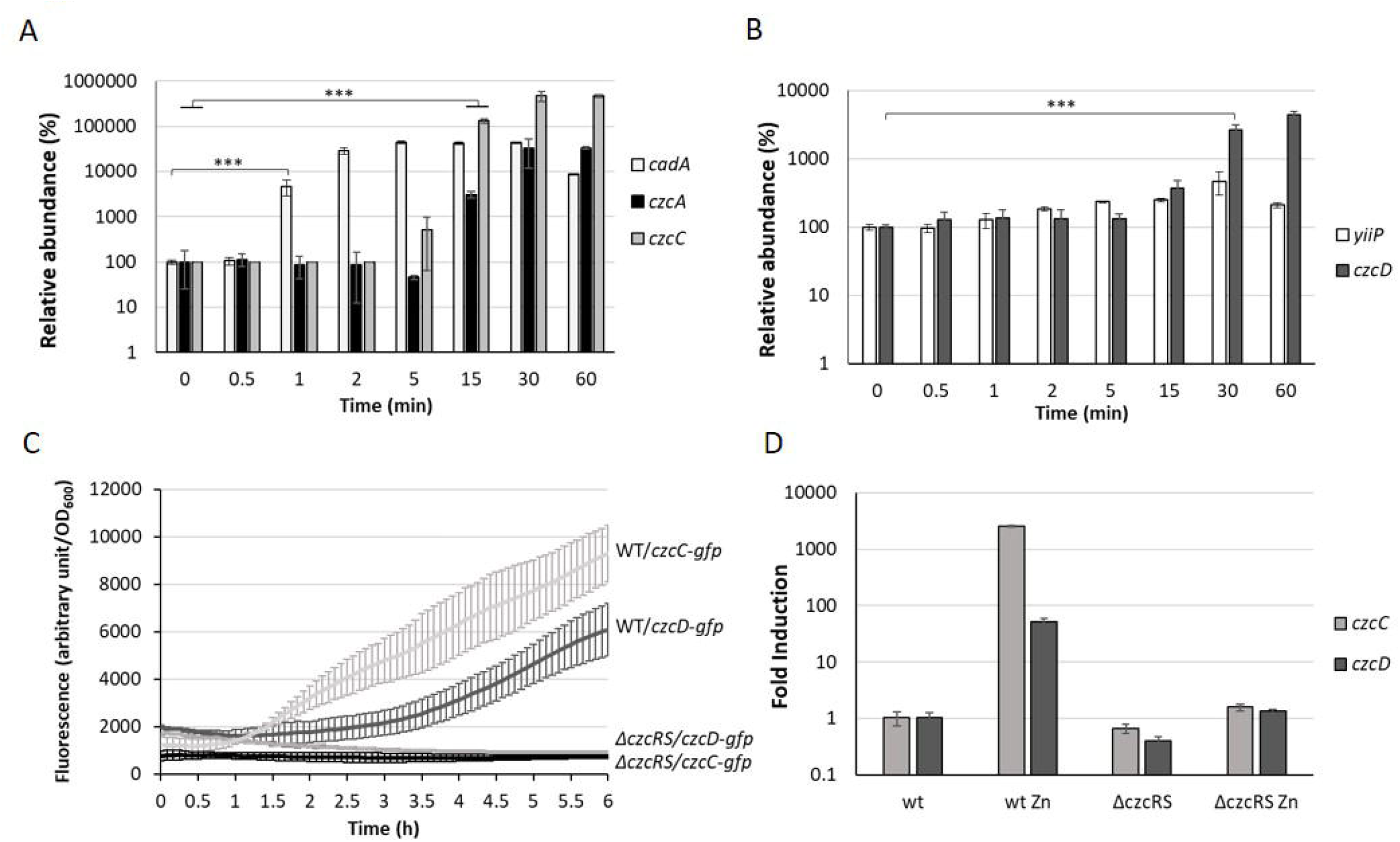
Fold expression of export systems. (A) *cadA* (P-type ATPase), *czcA* and *czcC* (RND) as well as (B) *yiip* and *czcD* (CDF) gene induction measured using the NanoString technology before and after addition of 2 mM ZnCl_2_ and monitored over the time. The 100% value corresponds to the number of mRNAs detected at t0 (before addition of Zn). Mean values of fold change compared to t0 and standard deviations (error bars) of three independent experiments are represented. Statistical analyses were performed according to the Student’s t-test and *p* values are given as follows: ≤ 0.001 (***). (C) Fluorescence measurement of *pczcC::gfp* (*czcC*) and *pczcD::gfp* (*czcD*) in the WT strain and the Δ*czcRS* mutant after addition of 2 mM ZnCl_2_. Values are normalized by optical density (OD_600_). Standard deviations (error bars) of three measurements are indicated. (D) qRT-PCR analysis of *czcC* and *czcD* mRNAs taken 2h after addition of 2 mM ZnCl_2_. The amount of mRNA is represented relative to time 0 (set to 1), before addition of ZnCl_2_. Results are normalized using the *oprF* gene and standard deviations (error bars) of three independent experiments are indicated.

After only 1 min, the amount of *cadA* mRNA rose to more than 20’000 transcripts, corresponding to a 50-fold induction (5000%, Fig 2A). The quantity of mRNA increased up to 5 min, reached a plateau, then decreased after 60 min (Fig 2A). Interestingly, a basal level of *cadA* was detected even under Zn-depleted conditions (t0) with more than 400 transcripts per 50 ng of total RNA (Table S3). The levels of *czcA* and *czcC* transcripts (last and first gene of the *czcCBA* operon) reached their maximum after 30 min. After 1 hour in the presence of the metal, a fold induction higher than 330 for *czcA* and higher than 4’700 for *czcC* was observed. Looking at the number of transcripts (Table S3), we noticed that, unlike *czcC, czcA* was already present under conditions of Zn limitation. Moreover, the number of *czcA* mRNA copies detected after 1 h in the presence of the metal was twice higher than that of *czcC*. Altogether, this suggested a transcriptional independence of *czcA* that could be mediated *via* an internal transcriptional start site (iTSS) within the *czcCBA* operon (30). A similar situation has been observed in the *czc* operon of *C. metallidurans*, giving rise to different polycistronic mRNAs, some constitutively expressed and additionally inducible in a CzcR-dependent manner (31).

### CzcD is part of the CzcRS regulon

CzcD and YiiP are two cation diffusion facilitators (CDF) that export Zn from the cytosol to the periplasm (26). Both showed low basal expression even before Zn addition (Table S3). However, the level of *yiip* mRNA did not increase upon Zn treatment, while *czcD* displayed a 26-fold induction, visible after 30 min (Fig 2B).

Since *czcD* is located on the *czc* locus in *C. metallidurans* (24), we wanted to determine whether *czcD* is part of the CzcRS-mediated regulation. For this purpose, we constructed a transcriptional fusion containing the *czcD* promoter controlling a *gfp* (green fluorescent protein) reporter gene. The expression of the fusion was tested in the wild type (WT) and Δ*czcRS* mutant strain in presence of 2mM of ZnCl_2_. Similarly to what is observed for the *czcC* promoter (12), we could show that *czcD* expression depends on the CzcRS TCS since no fluorescence was detected after Zn induction in the Δ*czcRS* mutant (Fig 2C). This result was also confirmed by qRT-PCR (Fig 2D). The fact that *czcD* gene induction is weaker than *czcCBA* (Table S3) might reflect its secondary role in Zn resistance (26,28). Altogether, these results suggest that *czcD*, although elsewhere in the genome from the *czcCBA* operon, is part of the CzcRS regulon.

### PA2807, a novel link between Zn and Cu homeostasis

In *C. metallidurans*, the *czc* element consists of nine genes with a *czcNICBADRSE* organization (31). We searched for conserved amino acid sequences in *P. aeruginosa* using DIAMOND (32). No significant CzcI and CzcN homologs were identified. Instead, at low stringency, we found that the PA2807 protein possesses two regions of approximately 40% sequence identity to *C. metallidurans* CzcE (Fig 3A). In *C. metallidurans*, CzcE is a periplasmic protein that has been shown to bind Cu but whose expression is also induced by Zn independently of the CzcDRS regulatory element (31,33). Using SignalP 5.0 software (34), a putative signal peptide of 48 amino acids that targets PA2807 to the periplasm is predicted with a probability of 0.47 (Fig 3A). To confirm the prediction, we tested the localization of a C-terminal 6 His-tagged PA2807 protein expressed *in trans* in *P. aeruginosa* and we found that it was strongly enriched in the periplasmic fraction (Fig 3B). The amino acid comparison with CzcE from *C. metallidurans* showed that PA2807 possesses an additional sequence of approximately 70 amino acids between the two conserved regions that contains 7 His residues and may constitute a metal binding site (Fig 3A).

**Figure 3:**
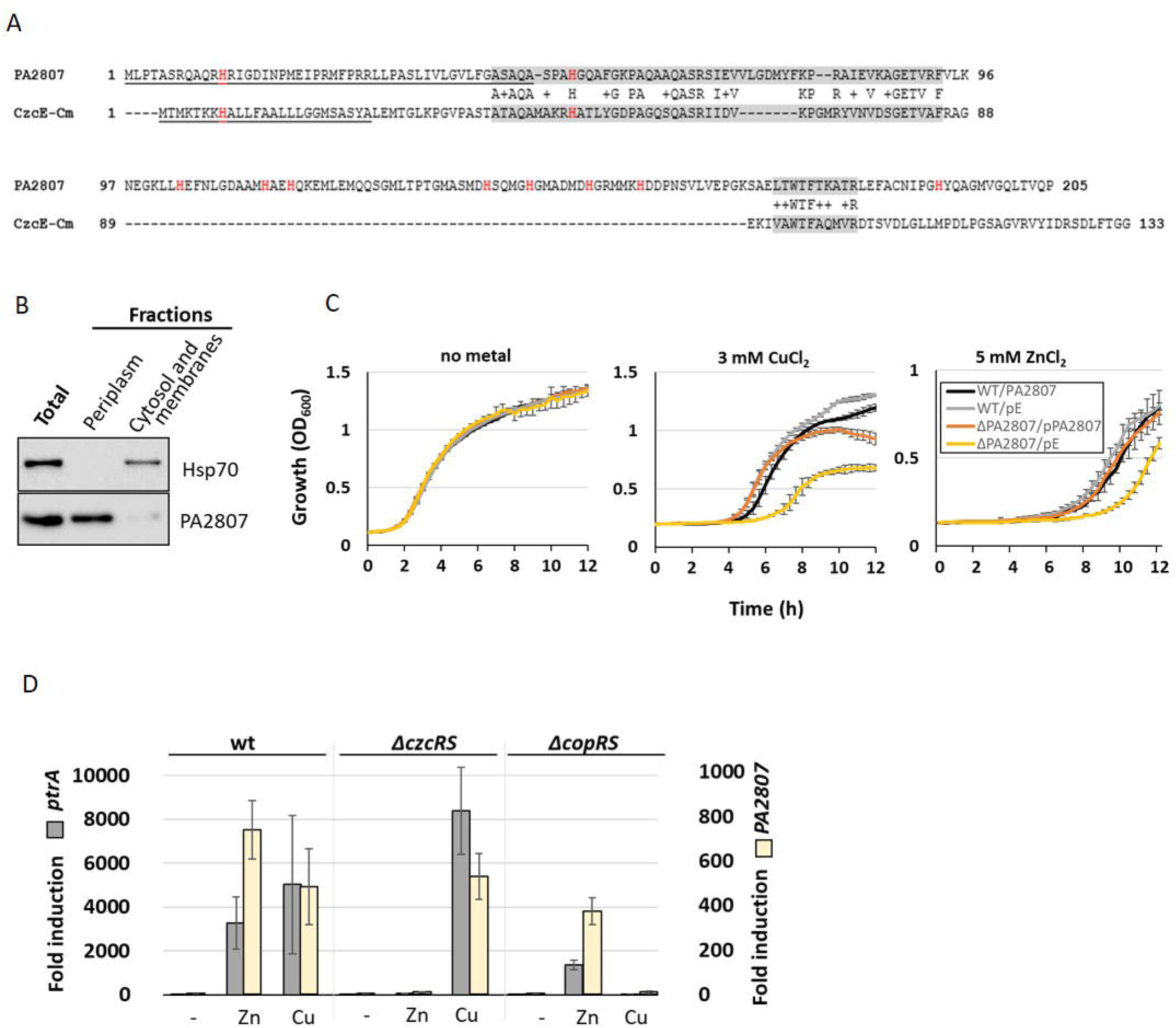
PA2807 encodes a CzcE-like protein. (A) Primary sequence alignment of PA2807 and CzcE from *Cupriavidus metallidurans*. The transit peptide is underscored and histidine residues are indicated in red. The two boxes of similarity are labelled in grey. (B) Immunoblot analysis of PA2807 localization. A culture of WT *P. aeruginosa* containing the 6His tagged version of the *PA2807* gene and its promoter on the pME6001 plasmid was induced for 2h with 2 mM CuCl_2_. 25 µg of total protein, and an equal volume (of 25 µg of protein) of periplasmic and cytosolic-membrane fractions was separated on a 4-12% SDS PAGE. Blots were exposed to anti-6His and anti-Hsp70 (loading and cytosolic control) antibodies. (C) Growth curves of WT *P. aeruginosa* and the Δ*PA2807* mutant carrying the empty IPTG-inducible pMMB66EH plasmid or the pMMB66EH-*PA2807* plasmid. Strains were cultivated in LB medium in the absence or presence of 3 mM CuCl_2_ or 5 mM ZnCl_2_ with 0.1 mM IPTG. (D) Fold induction of *ptrA* and *PA2807* mRNA analyzed using qRT-PCR on RNA extracted after 2h of growth in presence of 2 mM ZnCl_2_ or 2 mM CuCl_2_ as indicated. Error bars represent the standard deviations of three independent determinations.

*P. aeruginosa* PA2807 is described as a protein of the cupredoxin family and has been shown to be involved in Cu resistance (35). No effect on Zn resistance, however, was observed in a disc assay (36). To assess whether in our experimental settings the PA2807 protein is involved in Zn and/or Cu tolerance, we deleted this gene in a *P. aeruginosa* PAO1 strain and monitored the growth of the Δ*PA2807* mutant in the presence of 3 mM CuCl_2_ or 5 mM ZnCl_2_, as compared to the wild type strain (Fig 3C). The growth of Δ*PA2807* mutant was clearly delayed, indicating that the protein is indeed contributing to both Cu and Zn resistance. The mutant growth deficiency could be complemented by expressing the *PA2807* gene on a plasmid (Fig 3C). The involvement of PA2807 in Cu and Zn resistance was also confirmed on plate, using serial dilutions spotted on LB plates containing 5 mM ZnCl_2_ or 3 mM CuCl_2_ (Fig S1).

To determine whether the transcription of *PA2807* could be induced by Zn, we measured its transcript levels via the NanoString analysis. Surprisingly, it turned out to be the most induced gene, almost 10’000 times after 1h of induction (Table S3). *PA2807* is part of a gene cluster comprised of *ptrA* (*PA2806*), *PA2807* and *queF* (*PA2806*) next to the CopRS TCS (PA2809-PA2810) (35). PtrA is a small periplasmic protein involved in Cu resistance that has been shown to be also strongly induced in the presence of Zn (37,38). Using semi-quantitative RT-PCR, we found that the *PA2807* gene is co-transcribed as an operon with *ptrA* (Fig S2). Interestingly, the *ptrA-PA2807* transcript expression is not only induced by CopRS in the presence of Cu, but also in a CzcRS-dependent manner in presence of Zn, as shown by qRT-PCR (Fig 3D).

Altogether, these results demonstrated that the PA2807 protein of *P. aeruginosa* shares several characteristics with *C. metallidurans* CzcE and appears to be part of the CzcRS regulon in the presence of Zn.

### Hierarchy in the expression of export systems

The induction of the expression of these four Zn export systems, namely the CadA P-type ATPase, The RND CzcCBA, the two CDFs CzcD and Yiip, along with the newly discovered CzcE homolog, PA2807, showed a precise chronology of induction, as represented in Fig S3. Among all the export systems tested, *cadA* was the first to be induced (1 minute, early induced gene). The rapid induction is probably due to the CadR regulator already bound to *cadA* promoter region and therefore ready to transcribe *cadA* in the event of Zn excess (28). The *czc* regulon containing the *czcCBA* efflux pump, *czcD* and *PA280*7 (*czcE*-like) arrived later (15-30 min, late induced genes), while the gene encoding the CDF *yiiP* showed no induction under the conditions tested (uninduced gene).

## B) Zn uptake systems

Unlike the protein levels (Fig 1C), the mRNA levels of Zn import systems were strongly and rapidly repressed once Zn was added to the medium (Fig S3 and Table S3). Contrarily to export systems, however, no hierarchy in the regulation was evident and the decrease in the mRNA abundance of all uptake systems was statistically relevant (p value ≤ 0.001) between 2 to 5 minutes after Zn addition (Fig 4). One exception to this reduction was observed with the gene encoding the P-type ATPase HmtA, whose mRNA level remained constantly low (Table S3). This lack of repression could be due to the fact that this system is mainly involved in Cu import (39). Finally, among the three ABC transporters involved in Zn uptake, the PA4063-4066 system showed the highest expression, indicating that it could be the major player in Zn deficiency condition (Table S3).

**Figure 4:**
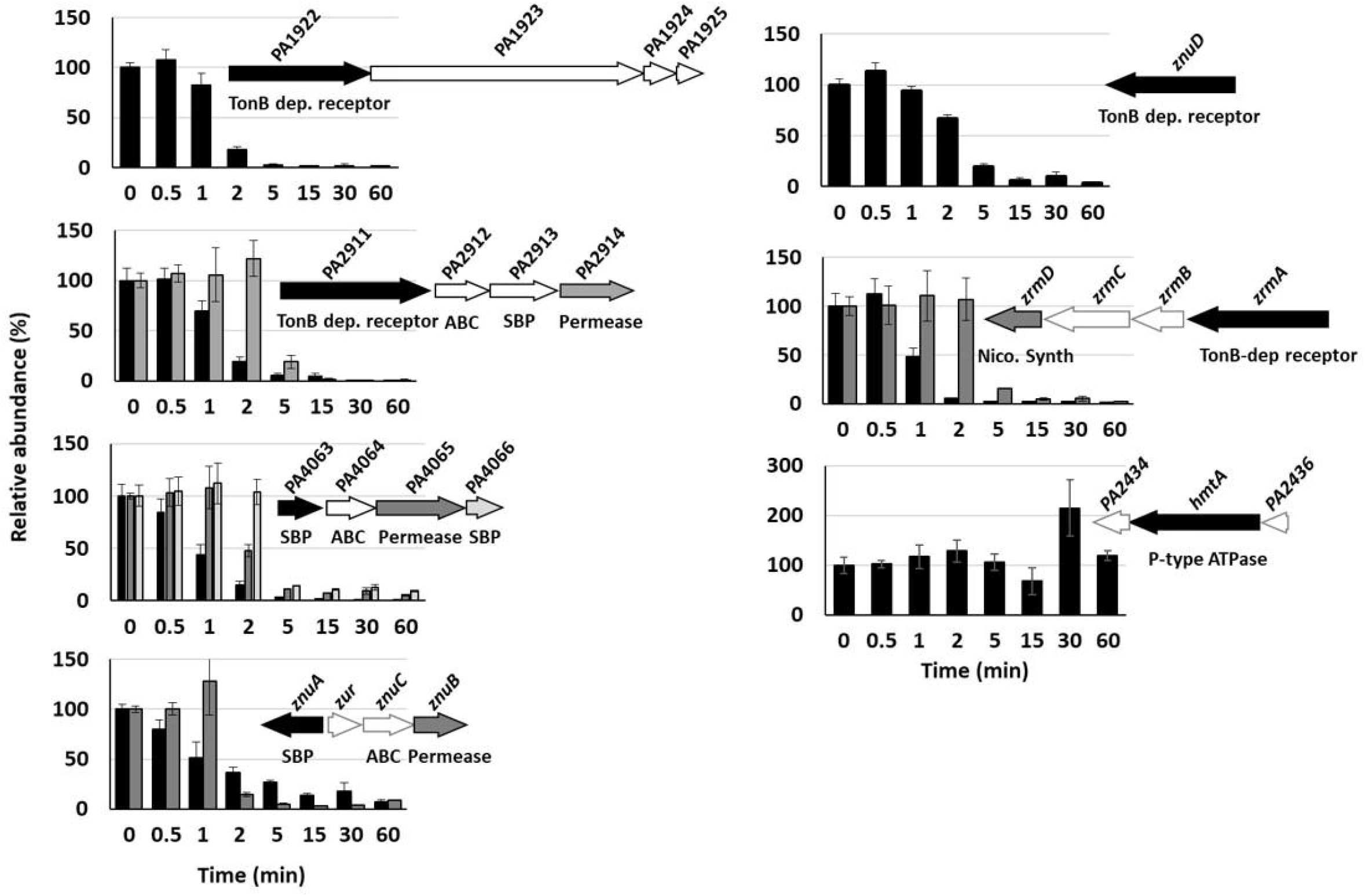
Repression of Zn import systems. The fold repressions over time of the seven import systems are shown. 100% corresponds to the number of mRNA copies at time 0 minute, immediately prior to addition of 2 mM ZnCl_2_. Black, dark and clear gray in the histogram correspond to the followed gene in the genomic arrangement represented next to the graph. White arrows represent genes of the system that were not followed during this work. The fold repression gives a p value ≤ 0.001 between 2 to 5 min after addition of Zn for all systems, except *hmtA* for which no repression was detected.

## C) Zn storage/release proteins

Under conditions of Zn deficiency, the stringent response regulator DksA is replaced by its DksA2 ortholog, which is devoid of a Zn finger domain (C-form) (21). When Zn reaches sufficient cytoplasmic concentration, *dksA2* transcription is switched off by the Zur protein and DksA takes over its place and function. In agreement with this, a rapid drop in the level of *dksA2* mRNA was observed: after 1 min of metal exposure only half of the number of mRNAs remained (Fig 5A and Table S3). A similar repression profile was shown for the two ribosomal genes *rpmE2* and *rpmJ*2 encoding Zn deficient orthologs of RpmE and RpmJ, respectively. Unlike their C-substitutes, the mRNA levels of *dksA, rpmE* and *rpmJ* remain constant, indicating the simultaneous expression of the three orthologous genes when the environment is depleted of metal. Are both C+ and C-proteins expressed at the same time or an additional regulatory mechanism occur at the next levels of gene expression? To address this question, we looked at the proteomic analysis focusing on the DksA and RpmE proteins (RpmJ was not detected) and their C-orthologs. (Fig 5B, Table S2). Intriguingly we observed that, while RpmE2 and DksA2 disappeared rapidly when Zn is added in excess, the proteins RpmE and DksA remain present, even in the absence of Zn.

**Figure 5:**
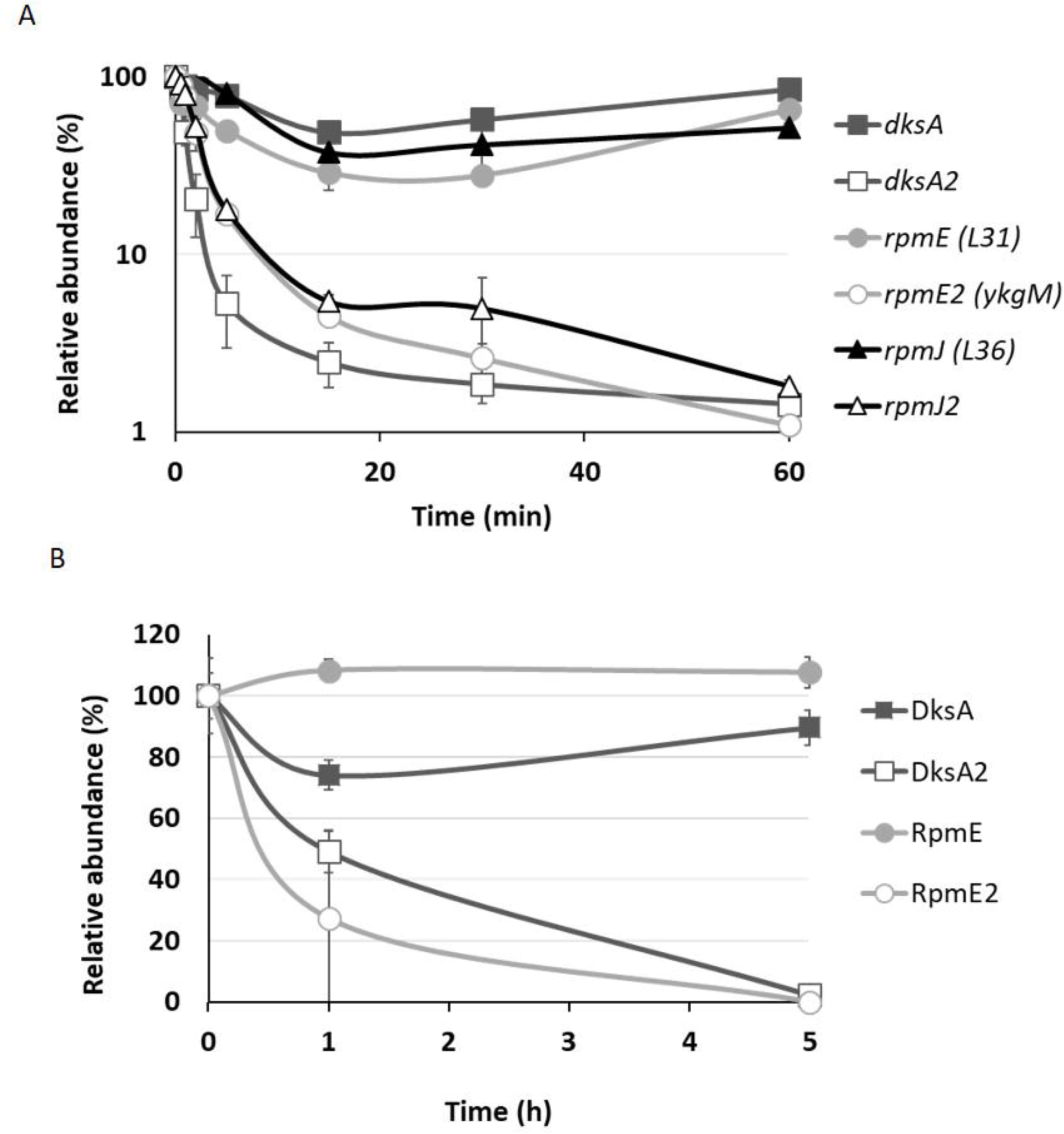
Kinetics of C_+_/C_-_ protein family. (A) Fold expression of *rpmE, rpmJ* and *dksA* genes and their C-paralogs over time, after addition of 2 mM ZnCl_2_. (B) Changes in the amount of DksA and RpmE proteins and their C-paralog after addition of 2 mM ZnCl_2_. 100% corresponds to the number of mRNA copies (A) or peptides (B) detected at t0. Mean values of fold change compared to t0 and standard deviations (error bars) of three independent experiments are represented.

## D) Zur affinity

The transcriptional repression of Zn import systems as well as the genes encoding C-proteins are mediated by the Zur protein. A Zur box was predicted for each of the repressed genes or operons (13) and Zur was shown to directly bind to the *znuA* and *dksA2* promoters (21,23). In *P. aeruginosa* this repressor is characterized by 2 Zn binding sites, the structural site located in the C-terminal region and a regulatory M-site (23). The Zur dimer contains 2 Zn^2+^ ions at the structural site and depending on the cytoplasmic Zn concentration, 0, 1 or 2 Zn^2+^ ions at the regulatory M-site. In *Bacillus subtilis*, promoter affinity depends on the amount of Zn linked to the regulatory site, allowing fine tuning of Zur regulation (40).

No clear hierarchy in the repression of Zur regulated genes could be observed with the NanoString experiment after addition of 2 mM ZnCl_2_ (Fig S3 and Table S3). We therefore used a more refined analysis to investigate a possible hierarchy in the affinity of the Zur regulator for the various putative Zur targets in *P. aeruginosa*. To this aim, we purified the Zur protein (Fig S4) and carried out Electrophoretic Mobility Shift Assay (EMSA) assays, in the absence (30 µM TPEN) or presence of Zn (5 µM ZnCl_2_) (Fig 6).

**Figure 6:**
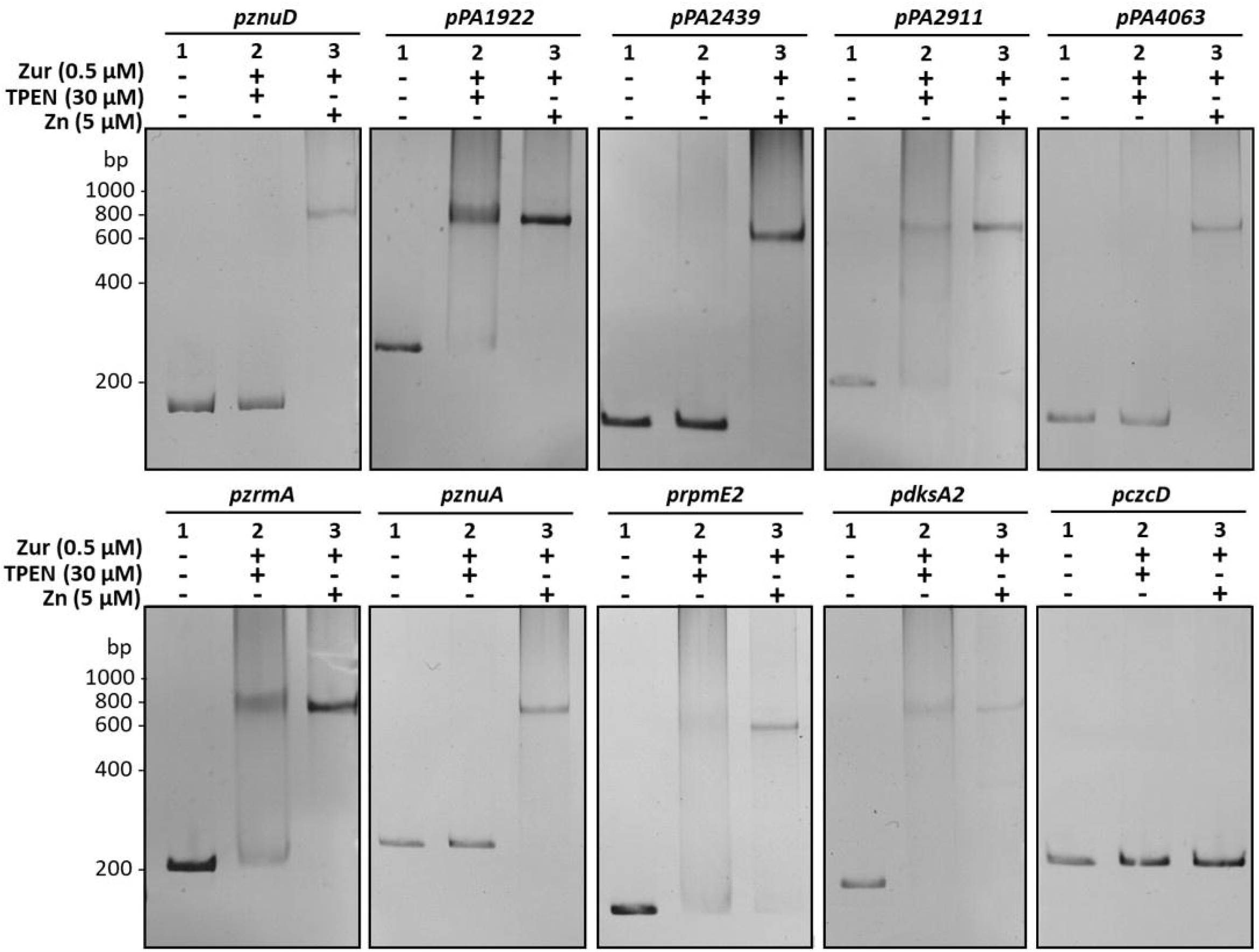
Zur binds to the diverse *zur* box with different affinities. Electrophoretic mobility shift assays using purified Zur protein and the different target promoters. The *czcD* promoter (*pczcD*) was used as a negative control. 30 ng of DNA were mixed with 1) 0 nM of Zur, 5 µM ZnCl_2_; 2) 500 nM Zur and 30 µM TPEN (Zn depleted condition) and 3) 500 nM Zur and 5 µM ZnCl_2_, as indicated. Reactions were loaded on non-denaturing 7.5% polyacrylamide gels, stained with ethidium bromide and viewed under UV light.

All the promoters tested were shifted, including the promoter of the PA2439 gene (first gene of the locus containing *hmtA*), for which a weak putative Zur binding site had previously been determined *in silico* (13). Interestingly, two distinct target profiles were emerging from the EMSA analysis. One set of targets exhibited a complete DNA-Zur interaction only in the presence of Zn (*znuA, znuD, PA4063 PA2439* and *rpmE2* promoters), while the other showed binding even in the absence of metal. This latter case represented the promoters of the *PA1922, PA2911* and *zrmA* genes, which encode for TonB-dependent receptor components (13) and the *dksA2* gene, involved in stringent response under Zn starvation conditions (21). Interestingly, these last four systems, are also involved in cobalt (Co) homeostasis, since *PA1922* is located within an operon that contains a *cobN*-like gene (*PA1923*), *PA2911* is co-transcribed with *PA2914*, a homolog of the cobalamin ABC permease (13) and pseudopaline, involving the *zrm* system, has been shown to transport metals such as Zn, Co, Fe, and nickel (Ni) (41). Similarly, *dksA2* is also part of an operon including *PA5535*, a *cobW*-like gene, involved in cobalamin biosynthesis. Altogether, these results reveal a putative Co-dependent regulatory activity of Zur, which could be linked with vitamin B12 (cobalamin) synthesis (42). Our EMSA conditions were not deficient in Co, hence, a sufficiently high concentration of this metal could be present to allow Zur-DNA interaction on such promoters, suggesting that Zur may be able to repress different targets on different promoters depending on the metal.

### Induction of carbapenem resistance

The response to Zn in *P. aeruginosa* is linked to carbapenem antibiotic resistance. The route of entry of these antibiotics into the cell is the OprD porin, whose expression is repressed in the presence of Zn at both transcriptional and post-transcriptional level via the CzcRS TCS and the Hfq RNA chaperone, respectively (12,43). Recently, we found that the metal concentrations present in phagolysosomes are sufficient to induce the Zn response and therefore carbapenem resistance in *P. aeruginosa* (28). We then followed the kinetics of carbapenem resistance induction by quantifying *oprD* downregulation at the mRNA and protein levels (Fig 7). A delay of 30 min was sufficient to observe a significant drop in *oprD* mRNA levels (Fig 7A) and the protein was only slightly present (less than 20%) after 5h of induction (Fig 7B, Table S1). This drop was also confirmed at protein levels by a Western blot experiment, where the protein was no longer detectable after 3h of metal addition (Fig 7C).

**Figure 7:**
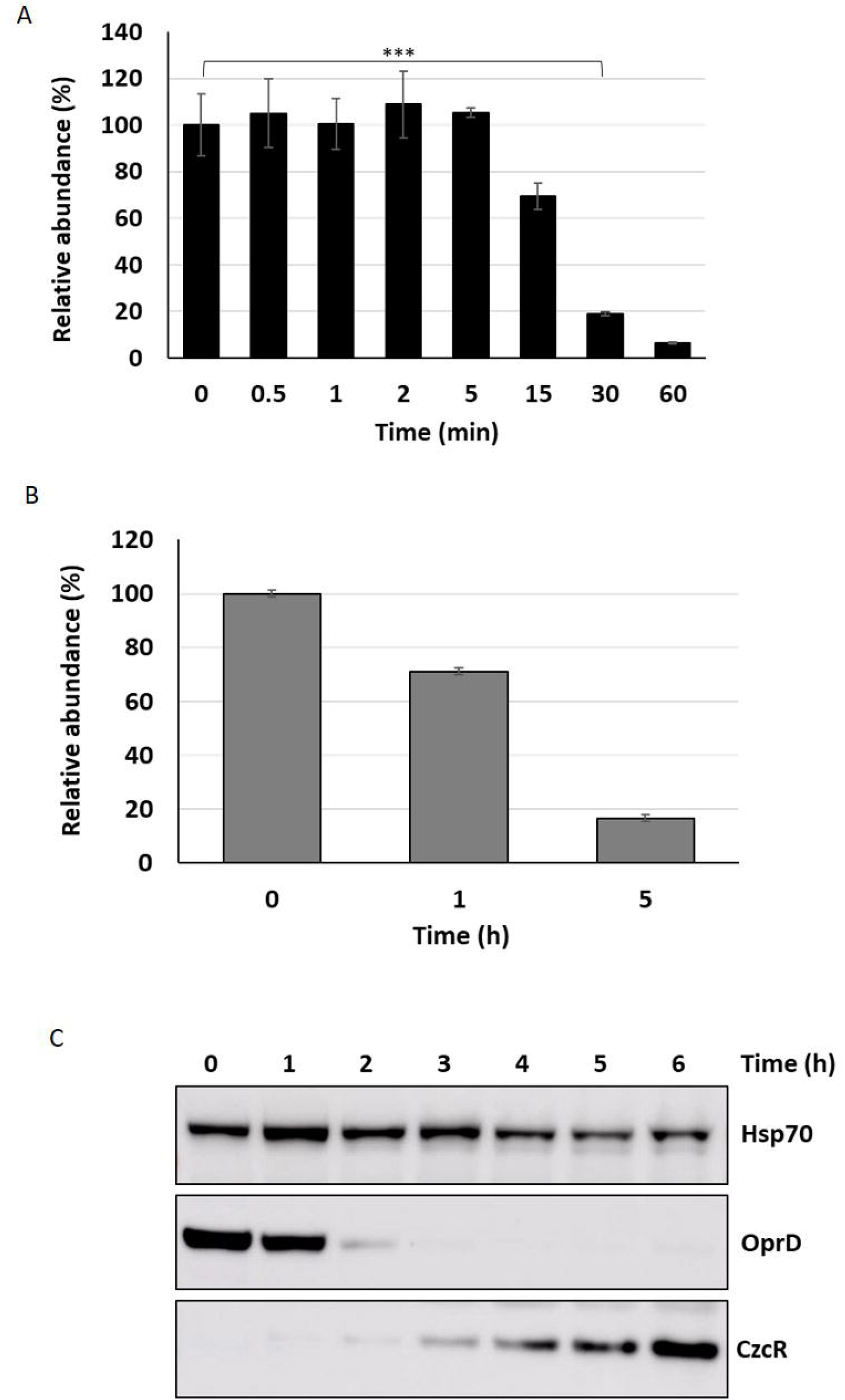
Proteomic and transcriptomic kinetics of OprD repression. (A) Percent decrease in *oprD* mRNA obtained by the NanoString technique after addition of 2 mM ZnCl_2_. 100% corresponds to the number of mRNA copies detected at t0. Mean values of fold change compared to t0 and standard deviations (error bars) of three independent experiments are represented. Statistical analyses were performed according to the Student’s t-test and *p* values are given as follows: ≤ 0.001 (***). (B) Decrease in OprD porin compared to t0 (100%), 1h and 5 h after addition of 2 mM Zn. (C) Western blot analysis of total protein sampled without Zn (t0) and after 1 to 6h following the addition of 2 mM ZnCl_2_. Blots were decorated with anti-OprD, anti-CzcR and anti-Hsp70 as loading control.

## Discussion

To infect and take advantage of its host, a pathogen must be able to adapt quickly to extreme conditions, including radical changes in Zn concentration. *P. aeruginosa* is typically a bacterium armed to counter this kind of situations, as shown by the numerous systems involved in Zn homeostasis. These systems, together with the ability to express them in a timely fashion, contribute to *P. aeruginosa* ability to infect hosts (10). In this study, we carried out a systematic investigation of the expression dynamics of the entire *P. aeruginosa* Zn homeostasis network when switching from a Zn depleted environment to a Zn excess situation, a transition mimicking what happens during an infectious process (44). Our analysis allowed us to draw a global model of Zn homeostasis gene expression organization in *P aeruginosa*, as illustrated in Fig 8.

**Figure 8:**
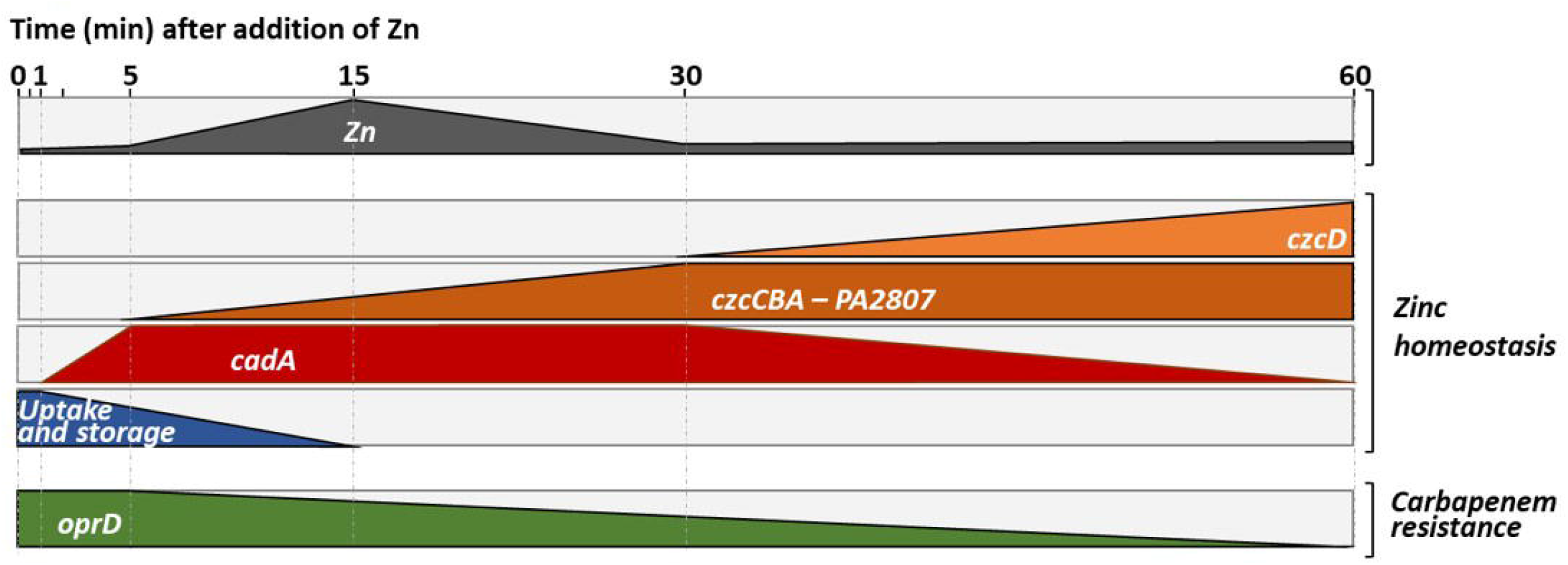
Representation of the dynamics of Zn homeostasis in *Pseudomonas aeruginosa*. The first block represents the intracellular Zn concentration, which increases when the cells move from a Zn-poor to a Zn-rich environment. An increase in intracellular Zn concentration is observed for up to 15 min, the time it takes for the import systems to be repressed and for the export system to start to eliminate the excess Zn (see second block and discussion). Later it appears that the ATP-consuming CadA system, is replaced by a less energy-demanding CzcD system that takes over to pump cytoplasmic Zn into the periplasm. This representation, highlights the repression of the expression of the OprD porin whose abundance is inversely proportional to resistance to the family of carbapenem antibiotics (last block).

As a first reaction, we observed a quick induction of *cadA* expression (after 1 min). Presumably, this serves to rapidly evacuate the Zn excess out of the cytoplasm into the periplasm (28). Downregulation of transcripts for uptake systems is also occurring fast, probably to prevent new Zn ions from entering the cell. Surprisingly, at the protein level this repression is visible only after 5 hours, suggesting a slow turnover of Zn uptake proteins. Given the importance of Zn for survival, this might be the result of a “prudential regulatory strategy”, to assure a prompt Zn uptake in case of the metal concentration decreases suddenly.

In a second step, the CzcCBA major efflux system is induced to detoxify all the cellular compartments, thus allowing the bacterium to thrive in an environment highly contaminated with Zn. In agreement with this, a decrease in total cellular Zn is observed as the expression of *czcCBA* increases (Fig 1D). The CzcD CDF is not an essential component in metal resistance (28), but we show here to be part of the *czcRS* regulon (Fig 2) and therefore responding to the periplasmic Zn concentration. Considering its late induction, CzcD may contribute to the maintenance of a steady state level of Zn resistance by continuing to pump Zn into the periplasm and allowing the CzcS to remain active while *cadA* expression decreases. One may consider that the P-type ATPase system CadA, despite being very efficient, uses ATP as an energy source while CzcD is driven by proton-motive forces and, therefore, might be more profitable for the cell in terms of energy optimization.

The *PA2807* gene was previously shown to be induced by the CopRS system in the presence of Cu (35). Here, we show that it encodes a CzcE-like protein involved in Zn resistance and also induced by the CzcRS systems in the presence of Zn. The role of this periplasmic protein in Zn and Cu resistance requires further investigation, but we could confirm the link between Zn and Cu resistance that has already been observed in other bacteria. For instance, in *Pseudomonas stutzeri*, a common overlapping response to Cu and Zn by common regulator DNA binding motifs has been elegantly demonstrated (45). In *P. aeruginosa*, the induction of CopRS induces the expression of the CzcRS TCS (46). In general, both Zn and Cu resistance may allow bacteria to resist to the toxic boost of metals that are discharged into the phagosome of macrophages to kill the pathogens (44,47).

Shift assays have shown a greater affinity of Zur for promoters controlling the expression of operons involved in cobalamin synthesis and therefore related to Co. Our *in vitro* conditions, by creating a Zn deficiency using TPEN, could displace the filling of Zur with Co (48). Some promoters would thus also appear to react first to a Co boost. The analysis of the sequences of the different Zur boxes did not allow us to highlight any particular signature. The DNA-Zur-Co interaction would deserve further investigation.

In *E. coli*, each of the 50’000 ribosomes present in exponentially growing cell contains about 3 Zn^2+^ ions, resulting in the recruitment of 75% of all the intracellular Zn available (21). Under conditions of Zn deficiency, the ribosomal proteins RpmE and RpmJ, along with the transcription factor DksA, all of which contain Cys4 Zn-ribbon motifs (and are thus called C+) are replaced by their C-paralog proteins (RpmE2, RpmJ2 and DksA2), which lack the key Cys residues and do not require Zn for their function. This replacement frees Zn, which becomes available for other essential biological tasks (49). A similar mechanism is found also in *B. subtilis*, in which the RpmE protein is degraded under Zn depleted conditions (50). As expected we found a rapid drop of C-proteins upon Zn excess in *P. aeruginosa*. However, unlike *B. subtilis, P. aeruginosa* C+ proteins are constitutively present even in conditions of Zn deficiency (t0). It is possible that the presence of these proteins reflects a regulatory strategy that guarantees a pool of proteins rapidly functional as soon as Zn is available.

In summary, this work shows for the first time the dynamics of the components of the Zn homeostasis network in *P. aeruginosa*. The use of the NanoString technology allowed us to precisely quantify several transcripts simultaneously and to follow their dynamics over time. The complementary proteomic analyses allowed us to observe the final outcome of this regulation and to visualize the Zn homeostasis systems at steady state (i.e. after 5h of growth in presence of Zn excess). Our findings are important to identify the molecular mechanisms favoring host colonization and infection, as well as to understand the environmental signals leading to the insurgence of antibiotic resistance. Specifically, the CzcRS-dependent repression of the *oprD* porin renders *P. aeruginosa* resistant to carbapenem in presence of Zn (12). According to our data, the OprD protein is no more detectable by western blot after 3h of Zn treatment (Fig 7C). We have recently reported OrpD repression during phagocytosis (28). Macrophages, and in general the immune system response, could therefore represent an important underestimated cause of carbapenem phenotypic resistance. Understanding the molecular mechanism favoring the appearance of carbapenem resistance will be thus fundamental to optimize the use of these last resort antibiotics.

## Material & Methods

### Bacterial Strains and Culture Media

The bacterial strains and plasmids used for this study are listed in Table S4. Cultures requiring initial Zn deficiency conditions were carried out in modified Luria-Bertani medium (AppliChem) supplemented with 30 µM of *N,N,N*′,*N*′-tetrakis(2-pyridinylmethyl)-1,2-ethanediamine (TPEN, Biotum), as described previously (28). Otherwise standard Luria-Broth medium (AppliChem) was used. Cultures were incubated at 37°C and when required, antibiotics were added to the medium at the following concentrations: 200 μg/mL carbenicillin (Cb, phytotechlab), 50 μg/mL Gentamycin (Gm, AppliChem) and 50 μg/mL tetracycline (Tc, Axxora) for *P. aeruginosa* or 100 μg/mL ampicillin (Ap, AppliChem) and 15 μg/mL Tc or Gm for *E. coli*.

### Genetic manipulations

DNA cloning was performed according to standard procedures (51). For Polymerase Chain Reactions, *P. aeruginosa* genomic DNA was used as template and the primers are listed in Table S5. Restriction and ligation enzymes (Promega) were employed according to the manufacturer’s instructions. Resulting recombinant plasmids were inserted into *E. coli* DH5α and confirmed by sequencing prior to being transformed into the *P. aeruginosa* wild type or the indicated mutant strains by electroporation (52).

Mutants were created according to the following strategy: two fragments flanking either the *PA2807* or the *copRS* TCS locus were amplified by PCR, digested with EcoRI/KpnI for the first insert and KpnI/BamHI for the second insert and ligated into the corresponding sites of the pME3087 suicide vector (53). After sequencing verification, recombinant plasmids were transformed into *P. aeruginosa* wild type strain. Merodiploids were resolved as previously mentioned (54) and the specific chromosomal deletions were confirmed by PCR amplification and sequencing.

For complementation of the Δ*PA2807* mutant, the full *PA2807* gene was amplified by PCR, digested with EcoRI and BamHI restriction enzymes and cloned into the corresponding sites of the IPTG-inducible pMMB66EH vector (55). The *PA2807* gene was also cloned with a C-terminal 6His tag into the pME6001 promoter-less vector (56). To this aim, the full gene and its promoter region were obtained by PCR and inserted into the pME6001 plasmid between the BamHI and HindIII restriction sites. For *czcD::gfp* fusion, the *czcD* promoter was amplified by PCR, digested with the KpnI and BglII enzymes and then ligated in the corresponding sites of the pBRR1-*gfp* plasmid (57).

### RNA extraction

Total RNA was isolated from the *P. aeruginosa* wild type or mutant strains as formerly explained (28). Briefly, overnight cultures were diluted to an optical density at 600 nm (OD_600_) of 0.1 and grown for 2h30 in M-LB containing 30 µM TPEN. 0.5 mL of each culture was mixed with 1 mL of RNA Bacteria Protect Reagent (Qiagen) immediately after 2 mM ZnCl_2_ induction (t0) and after several time points as indicated in the different figures. Total RNAs were extracted with an RNeasy mini kit (Qiagen) according to the manufacturer’s instructions. 5 µg of total RNA were treated with RQ1 RNase-free DNAse (Promega) for 2h at 37°C, followed by phenol/chloroform extraction and ethanol precipitation.

### NanoString nCounter expression analysis

mRNA content was analyzed at the iGE3 Genomics Platform (Faculty of Medicine, University of Geneva). 50 ng of total RNA were hybridized with multiplexed NanoString probes listed in Table S2 and samples were processed according to the published procedure (29). Barcodes were counted for 490 fields of view per sample. Background correction was performed by subtracting the mean +2 standard deviations of the negative controls for each sample. Values < 1 were set to 1. Positive controls were used as quality assessment: the ratio between the highest and the lowest average of the positive controls among samples was below 3. Counts for target genes were then normalized with the geometric mean of the four reference targets: *fbp, ppiD and rpoD*, selected according to (58) and *oprF*, usually used in the lab to normalize RT-qPCR. The stability of these genes has been verified using the geNorm algorithm (59).

### Quantitative RT-PCR analysis

500 ng of total RNA was reverse-transcribed using random hexamer primers (Promega) and Improm-II reverse transcriptase (Promega) according to the supplier’s instructions. The reverse transcriptase was then heat-inactivated and the resulting cDNAs were diluted tenfold in water. Quantitative PCR was performed in technical duplicates, using SYBR Select Master Mix (Applied biosystem), according to the supplier’s instructions. Results were analyzed as previously described (60) and normalized with the *oprF* gene.

### Semi-quantitative PCR analysis

500 ng of total RNA from WT *P. aeruginosa* grown in LB, LB supplemented with 2 mM ZnCl_2_ or with 2 mM CuCl_2_ was reverse-transcribed as described above and diluted tenfold in water. 28 cycles of PCR amplification were performed on the cDNAs and on the corresponding RNA dilution (negative control) as well as on genomic DNA (positive control) using the primers described in Table S5. Amplicons were analyzed on a 2% agarose gel stained with ethidium bromide using a standard procedure.

### GFP-reporter fusions

GFP fusion assays were carried out as previously described (28). Precultures of wild type and Δ*czcRS* strains carrying the *pczcC::gfp* or *pczcD::gfp* constructions were diluted to an OD_600_ of 0.1, transferred to a 96-well black plate (Costar) and grown for 2.5 h with shaking before being induced with 2 mM ZnCl_2_ (t0). Fluorescence at 528 nm was measured every 15 min using a Microplate reader (Synergy HT, BioTek instrument) and normalized with the OD_600_ values.

### Growth tests

Growth experiments were undertaken to investigate the metal susceptibility of the Δ*PA2807* mutant strain compared to the wild type, whether complemented or not with the *PA2807* gene. Overnight cultures of the different strains were diluted to an OD_600_ of 0.05 in LB medium containing 200 µg/mL carbenicillin and 0.1 mM IPTG (isopropyl β-D-1-thiogalactopyranoside). For growth curve analysis (Fig 3C), cultures were supplemented or not with either 3 mM CuCl_2_, or 5 mM ZnCl_2_, transferred to 96-well plates (Costar) and incubated at 37°C with shaking. Absorbance at 600 nm was measured every 15 min using a Microplate reader (Synergy HT, BioTek instrument). For growth spot assays (Fig S1), cultures were grown for 2h in LB containing 200 µg/mL carbenicillin and 0.1 mM isopropyl β-D-1-thiogalactopyranoside (IPTG; Axon Lab), then diluted to 10^−7^. 10 µl of each dilution was spotted onto LB plates, supplemented or not with either 3 mM CuCl_2_ or 5 mM ZnCl_2_ and incubated at 37°C for 24h.

### Western blot analyses

For PA2807 cellular localization, a fractionation procedure was adapted from (37). For this purpose, an overnight culture of a WT strain containing the 6His-tagged PA2807 version was diluted to an OD_600_ of 0.1 in LB supplemented with 50 µg/mL gentamycin and 2 mM CuCl_2_. After 5h of growth, 1 mL of cells was harvested and total proteins were solubilized at 2 mg/mL in 2X SDS-gel sample buffer (51) (an OD_600_ of 1 gives 0.175 mg/mL of protein). In parallel, 15 mL of culture were washed once with TMP buffer (10 mM Tris-HCl, 200 mM MgCl_2_, 1 mM AEBSF, pH8). Cells were then resuspended in 1 mL TMP buffer containing 0.5 mg/mL lysozyme (Fluka) and incubated at room temperature for 30 min before spinning down. The supernatant corresponds to the periplasmic fraction. The pellet, corresponding to membrane plus cytosolic fractions, was resuspended in the same volume of TMP buffer and sonicated. Both fractions were mixed with an equal volume of 2X SDS-gel sample buffer.

In order to investigate the kinetics of OprD protein repression over the time, an overnight WT *P. aeruginosa* culture was diluted to an OD_600_ of 0.05 in M-LB supplemented with 30 µM TPEN and incubated for 2h30 at 37°C. 1 mL of culture was collected and centrifuged immediately prior to being induced with 2 mM ZnCl_2_ and each hour as indicated in Fig 7C. All pellets were solubilized in the appropriate volume of 2X SDS-gel sample buffer to a total protein concentration of 2 mg/mL (an OD_600_ of 1 gives 0.175 mg/mL protein). Proteins were separated on SDS PAGE, using 4-12% precast gels (Invitrogen). Membrane transfer was performed with an iBlot 2 transfer stack (Invitrogen), according to the manufacturer’s instructions. Nitrocellulose membrane was incubated with anti-OprD or anti-penta-His and anti-Hsp70 antibodies as previously described (11). Blots were revealed by chemiluminescence (SuperSignal, Thermofisher), using the Amersham Imager 680 System.

### Intracellular Zn concentration

The procedure was adapted from (61). An overnight culture was diluted to an OD_600_ of 0.1 in M-LB supplemented with 30 µM TPEN and incubated 2h30 at 37°C. 1 mL of cells were collected and washed once with Phosphate Buffered Saline (PBS, Gibco) containing 1 mM ethylenediaminetetraacetic acid (EDTA, Promega). The culture was then induced with 2 mM ZnCl_2_ and a 1 mL-sample was collected at 5, 15, 30 and 60 min after Zn addition and processed in the same manner, to remove any traces of Zn. Pellets from the whole kinetic sequence were deep-frozen, freeze-dried, and kept in the dark pending analyses. Total dry material was weighed and transferred into polypropylene (PP) tubes. Acid digestion was then performed for 3 h at 90°C placing the PP tubes on Teflon heating blocks after adding 2 mL hydrochloric acid and 1 mL nitric acid (10 M HCl and 14 M HNO_3_, respectively, both Suprapur, Merck) following protocol adapted from (62) for trace metal quantification in biological matrices. Cooled digestates were then diluted in 10 mL ultra-pure deionized water (Milli-Q, 18.2 MΩ.cm, Millipore) and centrifuged at 4,000 rpm for 10 min (20 °C). The supernatant was stored in acid-cleaned PP tubes. Total Zn concentrations were quantified by Inductively Coupled Plasma-Mass Spectrometry (ICP-MS, 7900 Agilent) in samples diluted 10 fold with 1 % HNO_3_. Since no Certified Reference Materials (CRMs) exist for trace metal content in bacteria; other certified biological matrices were quantified, consisting in plankton material and seaweed (BCR®-414 and CD®200, respectively). Their analyses provided satisfactory results with recoveries for Zn concentrations > 90% and precision of ∼ 10 % (n = 6). At least triplicate of each condition was analyzed (vertical error bars in the graphics). Detection limits of ∼ 0.15 µg/L (3 x blank standard deviation) were reached for Zn concentrations for typical bacteria cell number of ∼ 8.10^8^.

### Zur expression and purification

The *zur* gene was amplified by PCR, cut with BamHI and EcoRI restriction enzymes and cloned into the pGEX-2T vector. The resulting plasmid was then transformed into the *E. coli* BL21 strain. Induction and purification were performed as described previously for the CadR protein (28). After removal of the GST tag with the thrombin protease, the protein was dialyzed-concentrated against PBS containing 1 mM dithiothreitol (DTT) and 50% glycerol. Protein purity was checked on SDS-Page (Bio-Rad) stained with Coomassie blue (Fig S4) and stored at -70°C until use.

### Electrophoretic Mobility Shift Assay (EMSA)

For the EMSA experiments, all DNA promoters indicated in Figure 6 were obtained by PCR and purified on agarose gel. Binding assays were performed according to the procedure already described (28). Briefly, reactions were performed with a mixture composed of the 5X Zn-less Binding Buffer (50 mM Tris, 200 mM KCl, 50 mM MgCl_2_, 5 mM DTT and 25% Glycerol), 30 ng of DNA, a Zn excess (5 µM) or deficiency (30 µM TPEN) in presence or absence of 500 nM Zur protein and incubated at room temperature for 30 min. Samples were then separated at 4°C on a 7.5% polyacrylamide native gel containing 2.5% glycerol in Tris Borate Buffer. Binding capacity was analyzed by staining the gel with 0.1% ethidium bromide and revealed with UV light using a NuGenius instrument.

### Proteomic analyses

Expression of the DksA and RpmE C-, C+ and OprD proteins was investigated by label-free proteomic analysis. Briefly, an overnight culture of the WT *P. aeruginosa* strain was diluted to an OD_600_ of 0.1 in M-LB supplemented with 30 µM TPEN and incubated for 2h30 at 37°C. 1 mL of culture was collected immediately before (t0) and after 1h and 5h induction with 2 mM ZnCl_2_. A culture pellet was resuspended in an appropriate volume of 50 mM NH_4_CO_3_ buffer in order to adjust the concentration to 0.5 mg/mL total proteins. Cells were digested with 0.1% RapiGest SF Surfactant (Waters) for 30 min at 60°C followed by 5 min at 100°C.

Extract contents were analyzed by ElectroSpray Ionization-Liquid Chromatography– Mass/Mass Spectrometric (ESI-LC-MS/MS) at the Proteomics Core Facility (Faculty of Medicine, University of Geneva). Raw data were processed using Proteome Discoverer 2.3 Software. Label-free quantification was performed using the “Top 3 precursor Intensity” method. Normalization was applied as well as an ANOVA analysis of variance with Bonferroni multiple test correction. A minimum of 10^5^, corresponding to the minimum detection value, was fixed for targets with zero values.

### Experimental relevance and statistical data

All experiments were performed, at least, in triplicate. For tables and graph representations, mean values or fold changes are shown in the figures, along with the standard deviations. When indicated, statistical analysis was performed according the Student’s t-test and significance p value was set to p ≤ 0.001 (***). For others, the figures show an indicative experiment.

## Acknowledgements

This work was supported by funding from the Swiss National Science Foundation (Grant 31003A_179336). MV is supported by an SNSF Ambizione grant (PZ00P3_174063). ISC is funded by the Gerbert Rüf Stiftung (Basel, Switzerland), Project Microbials GRS-071/17. MA is supported by the Portuguese FCT (contract, CEECIND/01777/2018). We are thankful for the expertise and advice of Dr Mylene Docquier from the Genomics Platform as well as the Proteomics Core Facility team, both from the Faculty of Medicine of the University of Geneva, Switzerland. We greatly acknowledge Gilbert Pfister and Beat Jermann from the Service de l’Ecologie de l’Eau (SECOE) of the Geneva Canton for providing access to ICP-MS facilities and for their analytical support, as well as Dr Christel Hassler, Dr Isabelle Worms and Thomas Cherubini for technical support and access to ICP-MS facilities of the UNIGE.

## Supplemental table legends

**Table S1:** Proteomic results for systems involved in Zn homeostasis before and after addition of 2 mM ZnCl_2_, as indicated. Mean values of three independent experiments and standard deviations are indicated. Undetected proteins are highlighted in grey.

**Table S2** (excel file): NanoString codeset details

**Table S3** (excel file) : mRNA count of the target genes involved in zinc homeostasis over time, after addition of 2 mM ZnCl_2_. Mean values and standard deviations of three independent experiments are represented. The software assigned an arbitrary value of 1 to undetected mRNAs (grey boxes).

**Table S4:** Strains and plasmids used in this study.

**Table S5:** Primers used in this study

## Figure legends

**Figure S1:**
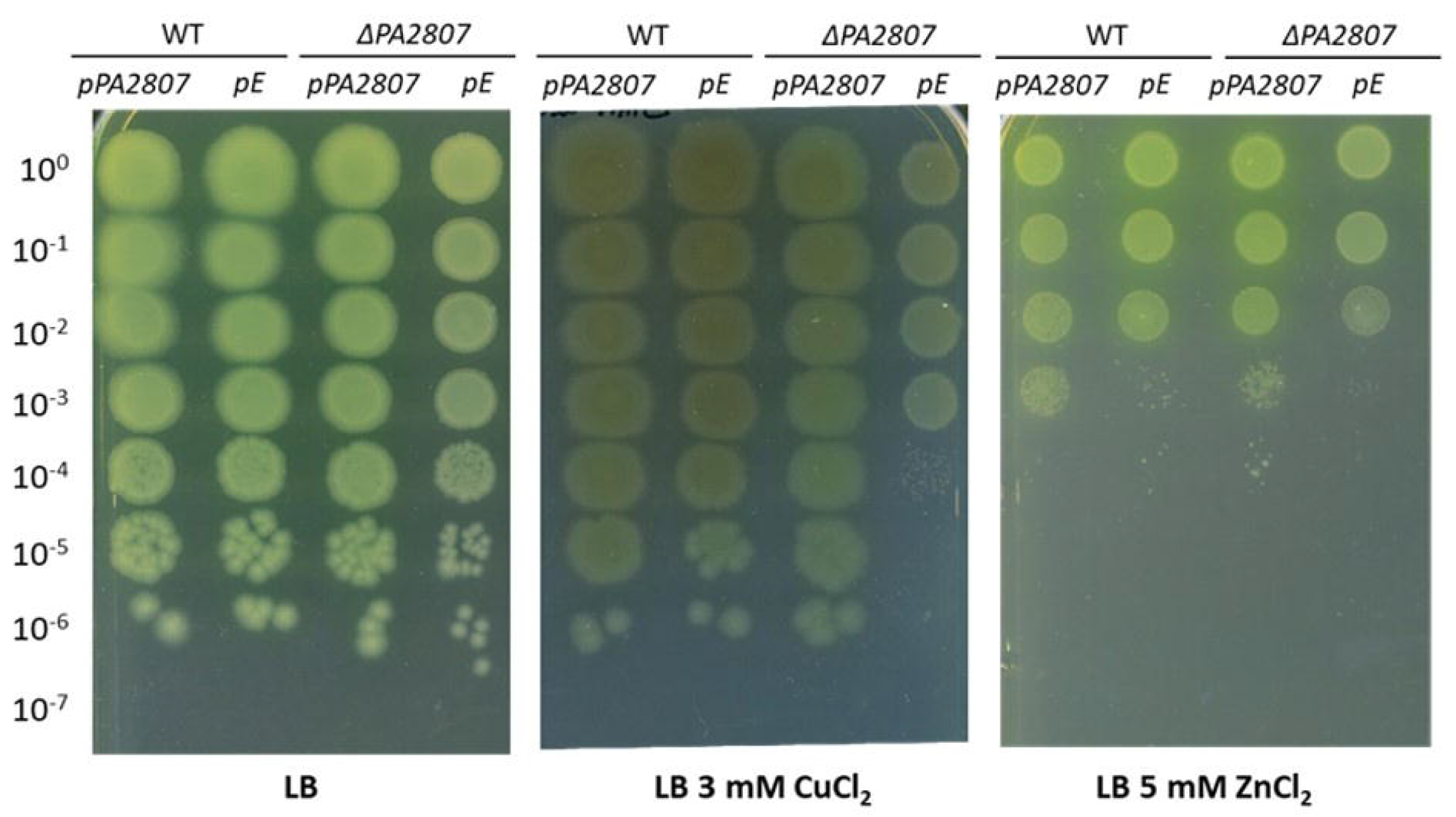
PA2807 and metal resistance. Spot test assay of a serial dilution of the WT and the Δ*PA2807* mutant carrying either an empty pMMB66EH plasmid (pE) or the *PA2807* gene cloned under the inducible *tac* promoter of the pMMB66EH plasmid (pPA2807). After induction with 0.1 mM IPTG 10 µL of the various dilutions were spotted onto LB, LB + 3 mM CuCl_2_ or 5 mM ZnCl_2_, as indicated and incubated for 24h at 37°C.

**Figure S2:**
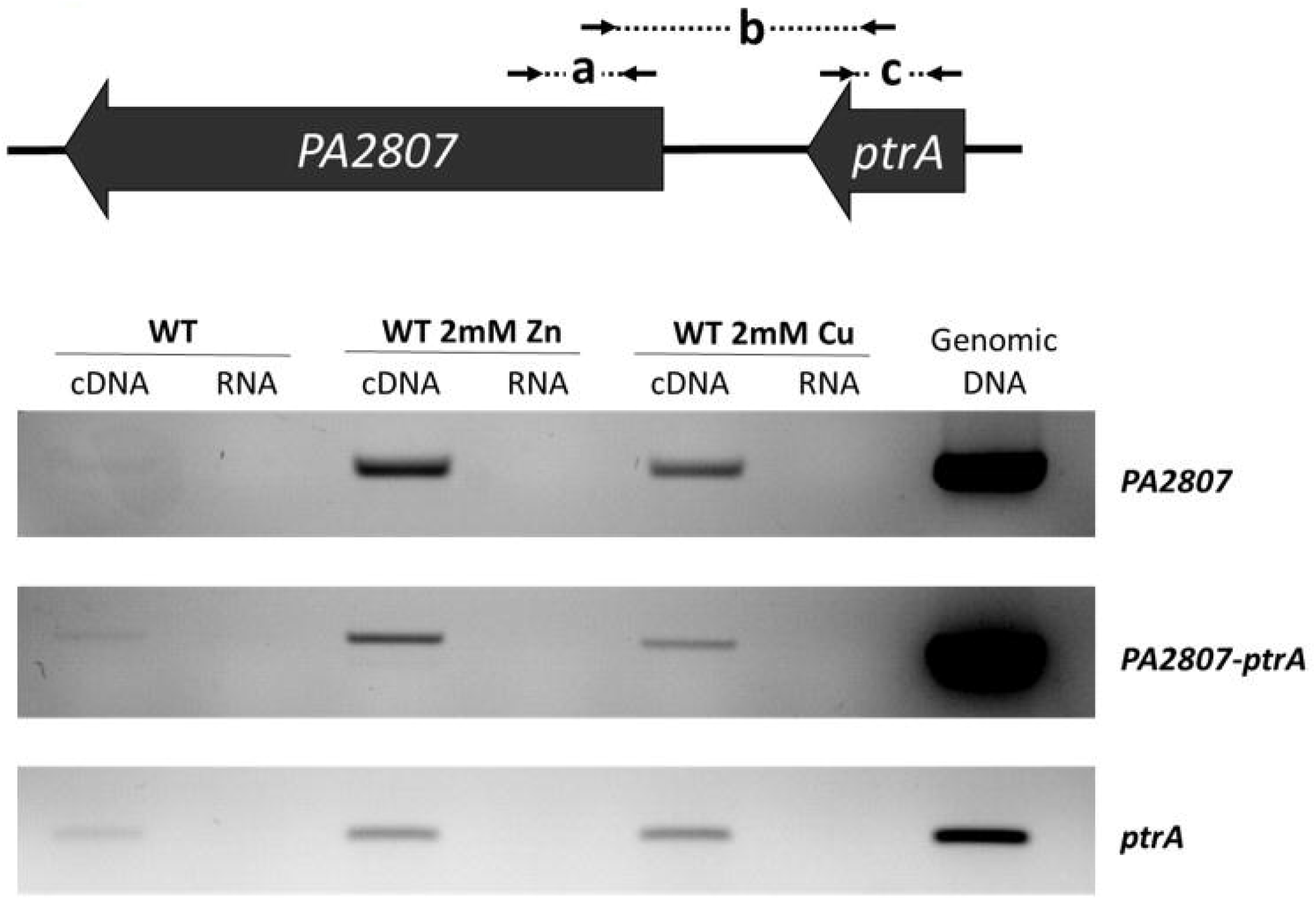
Co-transcription of the *ptrA* and *PA2807 genes*. Semi-quantitative RT-PCR analysis of the 3 amplicons (indicated in the upper panel) on RNA isolated from *P. aeruginosa* grown in LB, LB containing 2 mM ZnCl_2_ or 2 mM CuCl_2_. Primers used for each amplicon are indicated in the table S2. The amplification products were analyzed on a 2% agarose gel stained with ethidium bromide (lower panel). RNA corresponds to the negative control (without reverse transcriptase) and the Genomic DNA is used as positive control.

**Figure S3:**
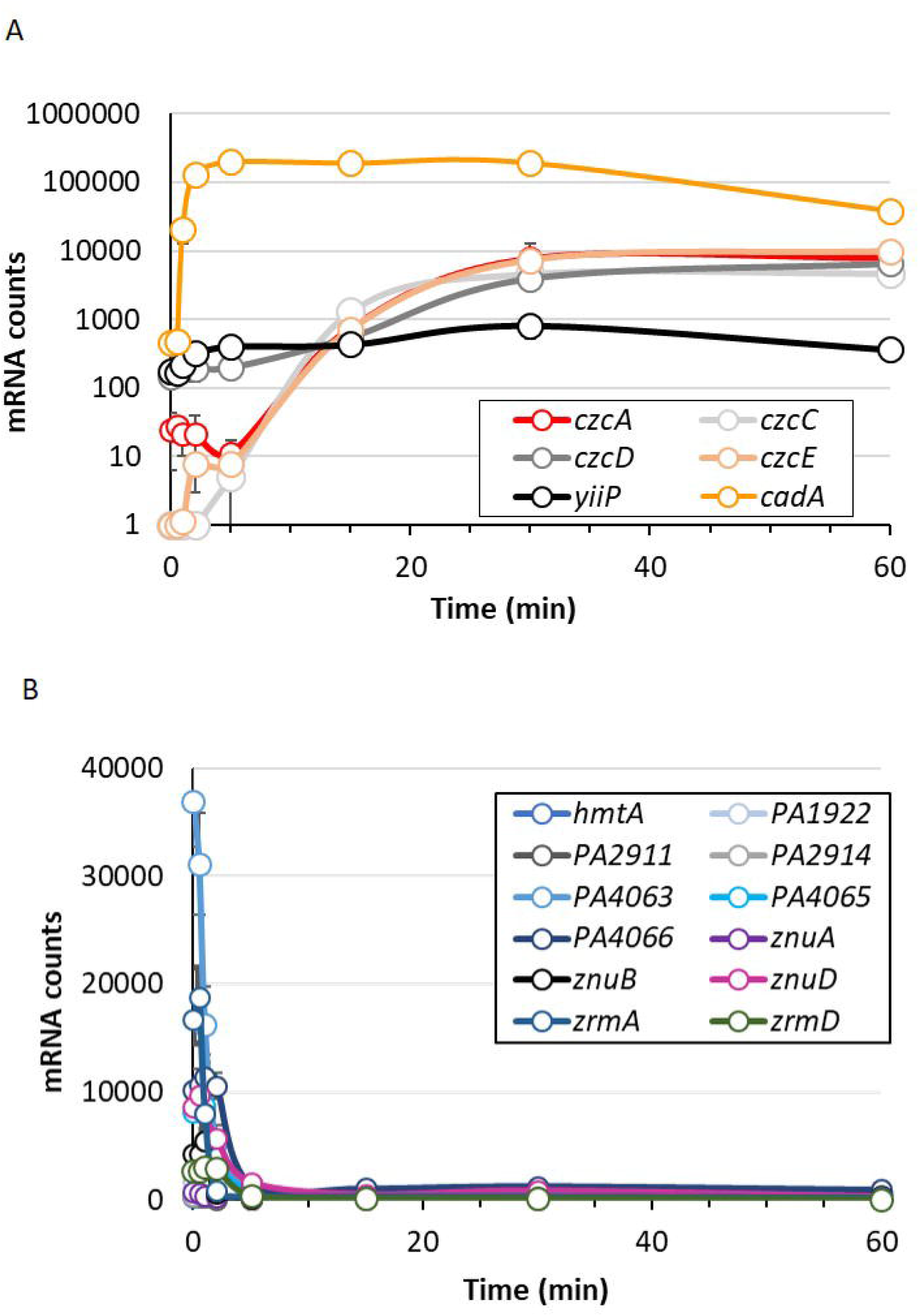
Transcription dynamics of genes involved in zinc homeostasis. mRNA counts of export (A) and import (C) systems as determined by NanoString analysis.

**Figure S4:**
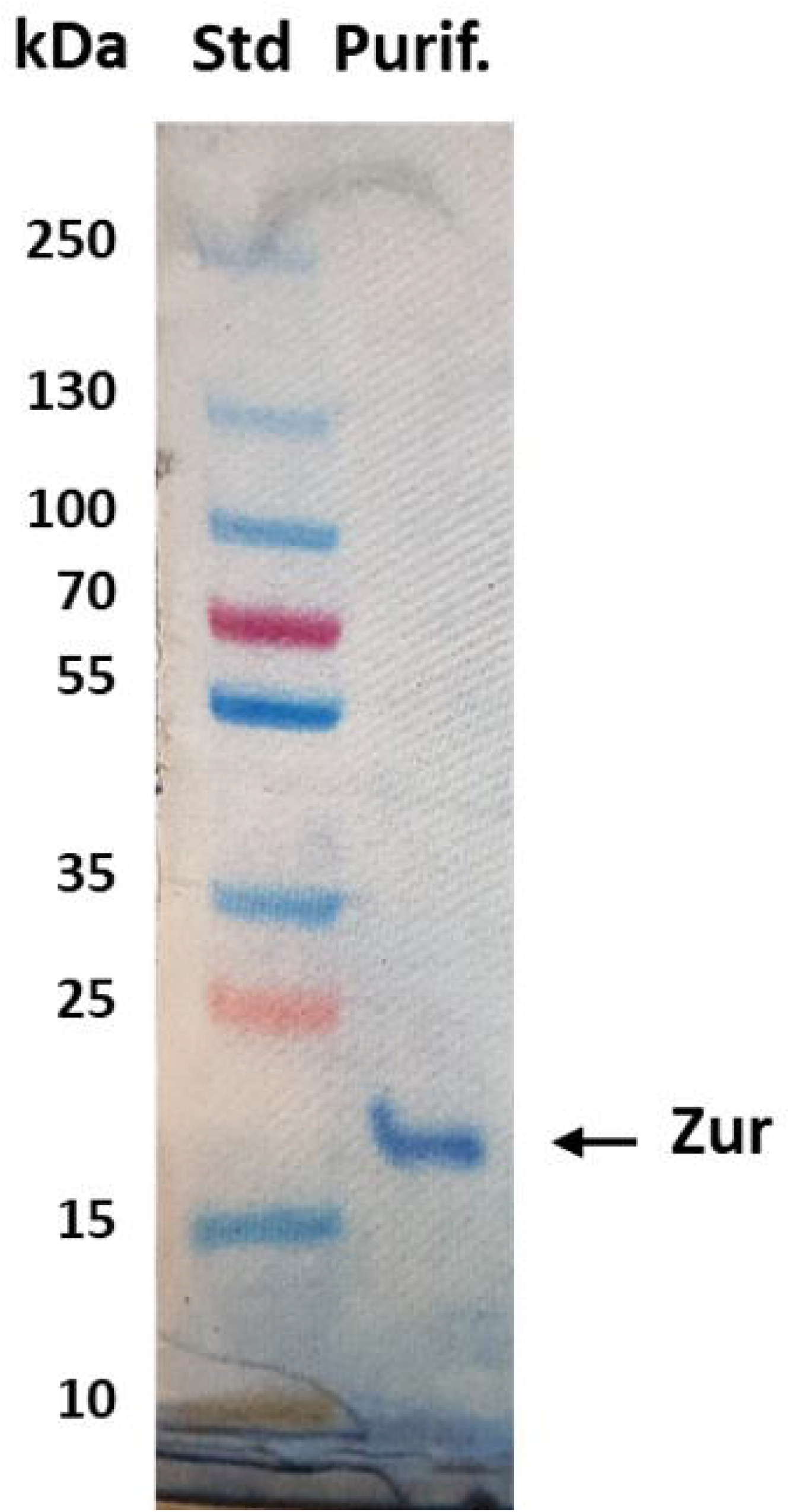
Purification of the Zur Protein. 5 µg of purified Zur protein were loaded onto a 4-12% SDS-PAGE and stained with Coomassie Blue.

## References

1. Blencowe, D.K. and Morby, A.P. (2003) Zn(II) metabolism in prokaryotes. FEMS Microbiol Rev, 27, 291–311.

2. Outten, C.E. and O’Halloran, T.V. (2001) Femtomolar sensitivity of metalloregulatory proteins controlling zinc homeostasis. Science, 292, 2488–2492.

3. Foster, A.W., Osman, D. and Robinson, N.J. (2014) Metal preferences and metallation. J Biol Chem, 289, 28095–28103.

4. Chandrangsu, P., Rensing, C. and Helmann, J.D. (2017) Metal homeostasis and resistance in bacteria. Nat Rev Microbiol, 15, 338–350.

5. Kehl-Fie, T.E. and Skaar, E.P. (2010) Nutritional immunity beyond iron: a role for manganese and zinc. Curr Opin Chem Biol, 14, 218–224.

6. Lonergan, Z.R. and Skaar, E.P. (2019) Nutrient Zinc at the Host-Pathogen Interface. Trends Biochem Sci, 44, 1041–1056.

7. Capdevila, D.A., Wang, J. and Giedroc, D.P. (2016) Bacterial Strategies to Maintain Zinc Metallostasis at the Host-Pathogen Interface. J Biol Chem, 291, 20858–20868.

8. Gao, H., Dai, W., Zhao, L., Min, J. and Wang, F. (2018) The Role of Zinc and Zinc Homeostasis in Macrophage Function. J Immunol Res, 2018, 6872621.

9. Kerr, K.G. and Snelling, A.M. (2009) Pseudomonas aeruginosa: a formidable and ever-present adversary. The Journal of hospital infection, 73, 338–344.

10. Gonzalez, M.R., Ducret, V., Leoni, S. and Perron, K. (2019) Pseudomonas aeruginosa zinc homeostasis: Key issues for an opportunistic pathogen. Biochim Biophys Acta Gene Regul Mech, 1862, 722–733.

11. Dieppois, G., Ducret, V., Caille, O. and Perron, K. (2012) The transcriptional regulator CzcR modulates antibiotic resistance and quorum sensing in Pseudomonas aeruginosa. PLoS One, 7, e38148.

12. Perron, K., Caille, O., Rossier, C., Van Delden, C., Dumas, J.L. and Kohler, T. (2004) CzcR-CzcS, a two-component system involved in heavy metal and carbapenem resistance in Pseudomonas aeruginosa. J Biol Chem, 279, 8761–8768.

13. Pederick, V.G., Eijkelkamp, B.A., Begg, S.L., Ween, M.P., McAllister, L.J., Paton, J.C. and McDevitt, C.A. (2015) ZnuA and zinc homeostasis in Pseudomonas aeruginosa. Sci Rep, 5, 13139.

14. Patzer, S.I. and Hantke, K. (1998) The ZnuABC high-affinity zinc uptake system and its regulator Zur in Escherichia coli. Mol Microbiol, 28, 1199–1210.

15. Mastropasqua, M.C., D’Orazio, M., Cerasi, M., Pacello, F., Gismondi, A., Canini, A., Canuti, L., Consalvo, A., Ciavardelli, D., Chirullo, B. et al. (2017) Growth of Pseudomonas aeruginosa in zinc poor environments is promoted by a nicotianamine-related metallophore. Mol Microbiol, 106, 543–561.

16. Lhospice, S., Gomez, N.O., Ouerdane, L., Brutesco, C., Ghssein, G., Hajjar, C., Liratni, A., Wang, S., Richaud, P., Bleves, S. et al. (2017) Pseudomonas aeruginosa zinc uptake in chelating environment is primarily mediated by the metallophore pseudopaline. Sci Rep, 7, 17132.

17. McFarlane, J.S. and Lamb, A.L. (2017) Biosynthesis of an Opine Metallophore by Pseudomonas aeruginosa. Biochemistry, 56, 5967–5971.

18. Hermansen, G.M.M., Hansen, M.L., Khademi, S.M.H. and Jelsbak, L. (2018) Intergenic evolution during host adaptation increases expression of the metallophore pseudopaline in Pseudomonas aeruginosa. Microbiology, 164, 1038–1047.

19. Haas, C.E., Rodionov, D.A., Kropat, J., Malasarn, D., Merchant, S.S. and de Crecy-Lagard, V. (2009) A subset of the diverse COG0523 family of putative metal chaperones is linked to zinc homeostasis in all kingdoms of life. BMC Genomics, 10, 470.

20. Ueta, M., Wada, C. and Wada, A. (2020) YkgM and YkgO maintain translation by replacing their paralogs, zinc-binding ribosomal proteins L31 and L36, with identical activities. Genes Cells.

21. Blaby-Haas, C.E., Furman, R., Rodionov, D.A., Artsimovitch, I. and de Crecy-Lagard, V. (2011) Role of a Zn-independent DksA in Zn homeostasis and stringent response. Mol Microbiol, 79, 700–715.

22. Furman, R., Biswas, T., Danhart, E.M., Foster, M.P., Tsodikov, O.V. and Artsimovitch, I. (2013) DksA2, a zinc-independent structural analog of the transcription factor DksA. FEBS Lett, 587, 614–619.

23. Ellison, M.L., Farrow, J.M., 3rd, Parrish, W., Danell, A.S. and Pesci, E.C. (2013) The transcriptional regulator Np20 is the zinc uptake regulator in Pseudomonas aeruginosa. PLoS One, 8, e75389.

24. Nies, D.H., Nies, A., Chu, L. and Silver, S. (1989) Expression and nucleotide sequence of a plasmid-determined divalent cation efflux system from Alcaligenes eutrophus. Proc Natl Acad Sci U S A, 86, 7351–7355.

25. Goldberg, M., Pribyl, T., Juhnke, S. and Nies, D.H. (1999) Energetics and topology of CzcA, a cation/proton antiporter of the resistance-nodulation-cell division protein family. J Biol Chem, 274, 26065–26070.

26. Salusso, A. and Raimunda, D. (2017) Defining the Roles of the Cation Diffusion Facilitators in Fe2+/Zn2+ Homeostasis and Establishment of Their Participation in Virulence in Pseudomonas aeruginosa. Front Cell Infect Microbiol, 7, 84.

27. Lee, S.W., Glickmann, E. and Cooksey, D.A. (2001) Chromosomal locus for cadmium resistance in Pseudomonas putida consisting of a cadmium-transporting ATPase and a MerR family response regulator. Appl Environ Microbiol, 67, 1437–1444.

28. Ducret, V., Gonzalez, M.R., Leoni, S., Valentini, M. and Perron, K. (2020) The CzcCBA Efflux System Requires the CadA P-Type ATPase for Timely Expression Upon Zinc Excess in Pseudomonas aeruginosa. Front Microbiol, 11, 911.

29. Geiss, G.K., Bumgarner, R.E., Birditt, B., Dahl, T., Dowidar, N., Dunaway, D.L., Fell, H.P., Ferree, S., George, R.D., Grogan, T. et al. (2008) Direct multiplexed measurement of gene expression with color-coded probe pairs. Nat Biotechnol, 26, 317–325.

30. Guell, M., Yus, E., Lluch-Senar, M. and Serrano, L. (2011) Bacterial transcriptomics: what is beyond the RNA horiz-ome? Nat Rev Microbiol, 9, 658–669.

31. Grosse, C., Anton, A., Hoffmann, T., Franke, S., Schleuder, G. and Nies, D.H. (2004) Identification of a regulatory pathway that controls the heavy-metal resistance system Czc via promoter czcNp in Ralstonia metallidurans. Arch Microbiol, 182, 109–118.

32. Buchfink, B., Xie, C. and Huson, D.H. (2015) Fast and sensitive protein alignment using DIAMOND. Nat Methods, 12, 59–60.

33. Zoropogui, A., Gambarelli, S. and Coves, J. (2008) CzcE from Cupriavidus metallidurans CH34 is a copper-binding protein. Biochem Biophys Res Commun, 365, 735–739.

34. Nielsen, H. (2017) Predicting Secretory Proteins with SignalP. Methods Mol Biol, 1611, 59–73.

35. Quintana, J., Novoa-Aponte, L. and Arguello, J.M. (2017) Copper homeostasis networks in the bacterium Pseudomonas aeruginosa. J Biol Chem, 292, 15691–15704.

36. Teitzel, G.M., Geddie, A., De Long, S.K., Kirisits, M.J., Whiteley, M. and Parsek, M.R. (2006) Survival and growth in the presence of elevated copper: transcriptional profiling of copper-stressed Pseudomonas aeruginosa. J Bacteriol, 188, 7242–7256.

37. Elsen, S., Ragno, M. and Attree, I. (2011) PtrA is a periplasmic protein involved in Cu tolerance in Pseudomonas aeruginosa. J Bacteriol, 193, 3376–3378.

38. Lei, L., Chen, J., Liao, W. and Liu, P. (2020) Determining the Different Mechanisms Used by Pseudomonas Species to Cope With Minimal Inhibitory Concentrations of Zinc via Comparative Transcriptomic Analyses. Front Microbiol, 11, 573857.

39. Lewinson, O., Lee, A.T. and Rees, D.C. (2009) A P-type ATPase importer that discriminates between essential and toxic transition metals. Proc Natl Acad Sci U S A, 106, 4677–4682.

40. Ma, Z., Gabriel, S.E. and Helmann, J.D. (2011) Sequential binding and sensing of Zn(II) by Bacillus subtilis Zur. Nucleic Acids Res, 39, 9130–9138.

41. Zhang, J., Zhao, T., Yang, R., Siridechakorn, I., Wang, S., Guo, Q., Bai, Y., Shen, H.C. and Lei, X. (2019) De novo synthesis, structural assignment and biological evaluation of pseudopaline, a metallophore produced by Pseudomonas aeruginosa. Chem Sci, 10, 6635–6641.

42. Osman, D., Cooke, A., Young, T.R., Deery, E., Robinson, N.J. and Warren, M.J. (2021) The requirement for cobalt in vitamin B12: A paradigm for protein metalation. Biochim Biophys Acta Mol Cell Res, 1868, 118896.

43. Ducret, V., Gonzalez, M.R., Scrignari, T. and Perron, K. (2016) OprD Repression upon Metal Treatment Requires the RNA Chaperone Hfq in Pseudomonas aeruginosa. Genes (Basel), 7.

44. Djoko, K.Y., Ong, C.L., Walker, M.J. and McEwan, A.G. (2015) The Role of Copper and Zinc Toxicity in Innate Immune Defense against Bacterial Pathogens. J Biol Chem, 290, 18954–18961.

45. Garber, M.E., Rajeev, L., Kazakov, A.E., Trinh, J., Masuno, D., Thompson, M.G., Kaplan, N., Luk, J., Novichkov, P.S. and Mukhopadhyay, A. (2018) Multiple signaling systems target a core set of transition metal homeostasis genes using similar binding motifs. Mol Microbiol, 107, 704–717.

46. Caille, O., Rossier, C. and Perron, K. (2007) A copper-activated two-component system interacts with zinc and imipenem resistance in Pseudomonas aeruginosa. J Bacteriol, 189, 4561–4568.

47. Botella, H., Stadthagen, G., Lugo-Villarino, G., de Chastellier, C. and Neyrolles, O. (2012) Metallobiology of host-pathogen interactions: an intoxicating new insight. Trends Microbiol, 20, 106–112.

48. Osman, D., Foster, A.W., Chen, J., Svedaite, K., Steed, J.W., Lurie-Luke, E., Huggins, T.G. and Robinson, N.J. (2017) Fine control of metal concentrations is necessary for cells to discern zinc from cobalt. Nat Commun, 8, 1884.

49. Gabriel, S.E. and Helmann, J.D. (2009) Contributions of Zur-controlled ribosomal proteins to growth under zinc starvation conditions. J Bacteriol, 191, 6116–6122.

50. Akanuma, G., Nanamiya, H., Natori, Y., Nomura, N. and Kawamura, F. (2006) Liberation of zinc-containing L31 (RpmE) from ribosomes by its paralogous gene product, YtiA, in Bacillus subtilis. J Bacteriol, 188, 2715–2720.

51. Sambrook, J., and D. W. Russell.. (2001.) Molecular cloning: a laboratory manual 3rd ed.Cold Spring Harbor Laboratory Press, Cold Spring Harbor, NY.

52. Choi, K.H., Kumar, A. and Schweizer, H.P. (2006) A 10-min method for preparation of highly electrocompetent Pseudomonas aeruginosa cells: application for DNA fragment transfer between chromosomes and plasmid transformation. J Microbiol Methods, 64, 391–397.

53. Voisard, C., Bull, C.T., Keel, C., Laville, J., Maurhofer, M., Schnider, U., Défago, G. and Haas, D. (1994), Molecular Ecology of Rhizosphere Microorganisms, pp. 67–89.

54. Ye, R.W., Haas, D., Ka, J.O., Krishnapillai, V., Zimmermann, A., Baird, C. and Tiedje, J.M. (1995) Anaerobic activation of the entire denitrification pathway in Pseudomonas aeruginosa requires Anr, an analog of Fnr. J Bacteriol, 177, 3606–3609.

55. Furste, J.P., Pansegrau, W., Frank, R., Blocker, H., Scholz, P., Bagdasarian, M. and Lanka, E. (1986) Molecular cloning of the plasmid RP4 primase region in a multi-host-range tacP expression vector. Gene, 48, 119–131.

56. Blumer, C., Heeb, S., Pessi, G. and Haas, D. (1999) Global GacA-steered control of cyanide and exoprotease production in Pseudomonas fluorescens involves specific ribosome binding sites. Proc Natl Acad Sci U S A, 96, 14073–14078.

57. Ouahrani-Bettache, S., Porte, F., Teyssier, J., Liautard, J.P. and Kohler, S. (1999) pBBR1-GFP: a broad-host-range vector for prokaryotic promoter studies. Biotechniques, 26, 620–622.

58. Gifford, A.H., Willger, S.D., Dolben, E.L., Moulton, L.A., Dorman, D.B., Bean, H., Hill, J.E., Hampton, T.H., Ashare, A. and Hogan, D.A. (2016) Use of a Multiplex Transcript Method for Analysis of Pseudomonas aeruginosa Gene Expression Profiles in the Cystic Fibrosis Lung. Infect Immun, 84, 2995–3006.

59. Vandesompele, J., De Preter, K., Pattyn, F., Poppe, B., Van Roy, N., De Paepe, A. and Speleman, F. (2002) Accurate normalization of real-time quantitative RT-PCR data by geometric averaging of multiple internal control genes. Genome Biol, 3, RESEARCH0034.

60. Schmittgen, T.D. and Livak, K.J. (2008) Analyzing real-time PCR data by the comparative C(T) method. Nature protocols, 3, 1101–1108.

61. D’Orazio, M., Mastropasqua, M.C., Cerasi, M., Pacello, F., Consalvo, A., Chirullo, B., Mortensen, B., Skaar, E.P., Ciavardelli, D., Pasquali, P. et al. (2015) The capability of Pseudomonas aeruginosa to recruit zinc under conditions of limited metal availability is affected by inactivation of the ZnuABC transporter. Metallomics, 7, 1023–1035.

62. Abdou, M., Gil-Díaz, T., Schäfer, J., Catrouillet, C., Bossy, C., Dutruch, L., Blanc, G., Cobelo-García, A., Massa, F., Castellano, M. et al. (2020) Short-term variations of platinum concentrations in contrasting coastal environments: The role of primary producers. Marine Chemistry, 222, 103782.

