## Supplementary material for "Global analysis of the zinc homeostasis network in *Pseudomonas aeruginosa* and its gene expression dynamics": Table S1

**Table S1:** Proteomic results for systems involved in Zn homeostasis before and after addition of 2 mM $ZnCl_2$, as indicated. Mean values of three independent experiments and standard deviations are indicated. Undetected proteins are highlighted in grey.

| | | | Average of AUC (Area Under the Curve) | | | | | | | | |
|---|---|---|---|---|---|---|---|---|---|---|---|
| Pathway | Protein | ID | t0 | | | 1h | | | 5h | | |
| Export | CzcA | PA2520 | **1.00E+05** | +/- | 0.00E+00 | **3.62E+08** | +/- | 4.71E+07 | **1.47E+09** | +/- | 4.51E+07 |
| | CzcB | PA2521 | **1.00E+05** | +/- | 0.00E+00 | **7.33E+08** | +/- | 2.36E+07 | **2.79E+09** | +/- | 9.50E+07 |
| | CzcC | PA2522 | **1.00E+05** | +/- | 0.00E+00 | **3.94E+08** | +/- | 3.60E+07 | **1.64E+09** | +/- | 9.29E+07 |
| | CadA | PA3690 | **1.93E+07** | +/- | 4.11E+06 | **1.18E+09** | +/- | 2.89E+07 | **4.42E+08** | +/- | 2.78E+07 |
| | CzcD | PA0397 | **1.00E+05** | +/- | 0.00E+00 | **6.67E+04** | +/- | 5.77E+04 | **2.38E+08** | +/- | 8.02E+06 |
| | YiiP | PA3963 | | | | | | | | | |
| Uptake | HmtA | PA2435 | **1.45E+07** | +/- | 6.86E+06 | **8.13E+06** | +/- | 0.00E+00 | **1.00E+05** | +/- | 0.00E+00 |
| | PA1922 | PA1922 | **2.27E+08** | +/- | 5.29E+06 | **1.49E+08** | +/- | 7.21E+06 | **1.64E+07** | +/- | 2.00E+06 |
| | PA1923 | PA1923 | **1.73E+08** | +/- | 3.92E+07 | **3.38E+07** | +/- | 7.53E+06 | **1.00E+05** | +/- | 0.00E+00 |
| | PA1924 | PA1924 | **1.60E+07** | +/- | 3.25E+06 | **1.00E+05** | +/- | 0.00E+00 | **1.00E+05** | +/- | 0.00E+00 |
| | PA1925 | PA1925 | **5.61E+07** | +/- | 1.31E+06 | **2.33E+07** | +/- | 3.21E+05 | **1.00E+05** | +/- | 0.00E+00 |
| | PA2911 | PA2911 | **3.57E+08** | +/- | 7.64E+06 | **2.35E+08** | +/- | 5.20E+06 | **5.79E+07** | +/- | 6.18E+06 |
| | PA2912 | PA2912 | **1.15E+08** | +/- | 3.21E+06 | **6.18E+07** | +/- | 1.39E+07 | **1.00E+05** | +/- | 0.00E+00 |
| | PA2913 | PA2913 | **2.06E+08** | +/- | 2.89E+06 | **1.35E+08** | +/- | 3.51E+06 | **8.34E+06** | +/- | 0.00E+00 |
| | PA2914 | PA2914 | | | | | | | | | |
| | PA4063 | PA4063 | **2.04E+09** | +/- | 6.56E+07 | **1.78E+09** | +/- | 4.93E+07 | **3.19E+08** | +/- | 1.85E+07 |
| | PA4064 | PA4064 | **2.68E+08** | +/- | 1.01E+07 | **1.69E+08** | +/- | 1.15E+07 | **2.26E+07** | +/- | 7.32E+06 |
| | PA4065 | PA4065 | **2.40E+08** | +/- | 2.23E+07 | **1.66E+08** | +/- | 7.94E+06 | **3.00E+07** | +/- | 5.20E+06 |
| | PA4066 | PA4066 | **1.30E+08** | +/- | 4.62E+06 | **9.66E+07** | +/- | 1.91E+06 | **1.00E+05** | +/- | 0.00E+00 |
| | ZnuA | PA5498 | **2.68E+08** | +/- | 2.01E+07 | **2.09E+08** | +/- | 8.19E+06 | **2.34E+07** | +/- | 1.12E+07 |
| | ZnuB | PA5501 | | | | | | | | | |
| | ZnuC | PA5500 | **6.94E+07** | +/- | 2.14E+07 | **7.68E+07** | +/- | 0.00E+00 | **1.00E+05** | +/- | 0.00E+00 |
| | ZnuD | PA0781 | **7.07E+09** | +/- | 3.67E+08 | **4.83E+09** | +/- | 2.86E+08 | **1.75E+09** | +/- | 1.39E+08 |
| | ZrmA | PA4837 | **8.61E+08** | +/- | 9.61E+06 | **5.35E+08** | +/- | 3.26E+07 | **1.06E+08** | +/- | 5.13E+06 |
| | ZrmB | PA4836 | **1.54E+08** | +/- | 5.03E+06 | **3.81E+07** | +/- | 5.83E+06 | **1.00E+05** | +/- | 0.00E+00 |
| | ZrmC | PA4835 | **9.13E+07** | +/- | 3.33E+06 | **6.00E+07** | +/- | 5.45E+06 | **1.00E+05** | +/- | 0.00E+00 |
| | ZrmD | PA4834 | **2.42E+07** | +/- | 0.00E+00 | **3.28E+07** | +/- | 0.00E+00 | **1.00E+05** | +/- | 0.00E+00 |
| Storage | DksA | PA4723 | **1.25E+09** | +/- | 4.00E+07 | **9.26E+08** | +/- | 6.17E+07 | **1.12E+09** | +/- | 7.21E+07 |
| | DksA2 | PA5536 | **6.85E+08** | +/- | 8.38E+07 | **3.36E+08** | +/- | 4.65E+07 | **1.52E+07** | +/- | 2.14E+06 |
| | RpmE | PA5049 | **2.09E+09** | +/- | 1.57E+08 | **2.26E+09** | +/- | 7.51E+07 | **2.25E+09** | +/- | 1.04E+08 |
| | RpmE2 | PA3601 | **2.48E+09** | +/- | 4.58E+07 | **6.77E+08** | +/- | 7.13E+08 | **1.00E+00** | +/- | 0.00E+00 |
| | RpmJ | PA4242 | | | | | | | | | |
| | RpmJ2 | PA3600 | | | | | | | | | |
| Others | CzcE | PA2807 | **1.00E+05** | +/- | 0.00E+00 | **1.00E+05** | +/- | 0.00E+00 | **1.46E+08** | **+/-** | 2.76E+07 |
| | OprD | PA0958 | **1.34E+09** | +/- | 1.53E+07 | **9.50E+08** | +/- | 1.64E+07 | **2.23E+08** | **+/-** | 1.55E+07 |
