## Supplementary material for "Global analysis of the zinc homeostasis network in *Pseudomonas aeruginosa* and its gene expression dynamics": Table S4

**Table S4:** Strains and plasmids used in this study.

| Strain or plasmid | Relevant characteristic(s)[a] | Reference or source |
|---|---|---|
| ***P. aeruginosa*** | | |
| Wild type | PAO1 wild type | laboratory collection |
| *ΔczcRΔczcS* | PAO1 Δ*czcR*Δ*czcS* | (1) |
| *ΔcopRΔcopS* | PAO1 Δ*copR*Δ*copS* | This study |
| *ΔPA2807* | PAO1 Δ*PA2807* | This study |
| ***E. coli*** | | |
| DH5α | recA1, endA1, hsdR17, deoR, thi-1, supE44, gyrA96, relA1, Δ(lacZYA-argF), U169(φ80dlacZΔM15) | (2) |
| BL21(DE3) | E. coli str. B F– ompT gal dcm lon hsdSB(rB–mB–) λ(DE3 [lacI lacUV5-T7p07 ind1 sam7 nin5]) [malB+]K-12(λS) | (3) |
| **Plasmids** | | |
| pME3087 | Suicide plasmid, Co1E1 replicon; $Tc^r$ | (4) |
| pMMB66EH | Expressing vector carrying an IPTG-inducible promoter; $Ap^r$, $Cb^r$ | (5) |
| pMMB66EH-*PA2807* | pMMB66EH derivative, carrying the *PA2807* gene; $Ap^r$, $Cb^r$ | This study |
| pME6001 | pME6000 derivative plasmid; $Gm^r$ | (6) |
| pME6001-*PA2807* | pME6001 carrying the PA2807 gene and its promoter; $Gm^r$ | This study |
| pBBR1-*gfp* | Transcriptional *gfp* fusion cloning vector; $Ap^r$, $Cb^r$ | (7) |
| *czcC:gfp* | pBBR1 derivative, carrying the *czcC* promoter; $Ap^r$, $Cb^r$ | (8) |
| *czcD:gfp* | pBBR1 derivative, carrying the *czcD* promoter; $Ap^r$, $Cb^r$ | This study |
| pGex-2T-*zur* | GST-fusion expression plasmid, carrying the *zur* gene; $Ap^r$ | This study |

[a]Antibiotic resistance is indicated by r: Ap ampicillin; Tc tetracycline; Cb carbenicillin; Gm gentamycin

### References:

1. Caille, O., Rossier, C. and Perron, K. (2007) A copper-activated two-component system interacts with zinc and imipenem resistance in Pseudomonas aeruginosa. *J Bacteriol*, **189**, 4561-4568.
2. Sambrook, J., and D. W. Russell. . (2001. ) *Molecular cloning: a laboratory manual* 3rd ed.Cold Spring Harbor Laboratory Press, Cold Spring Harbor, NY.
3. Studier, F.W., Rosenberg, A.H., Dunn, J.J. and Dubendorff, J.W. (1990) Use of T7 RNA polymerase to direct expression of cloned genes. *Methods in enzymology*, **185**, 60-89.
4. Voisard, C., Bull, C.T., Keel, C., Laville, J., Maurhofer, M., Schnider, U., Défago, G. and Haas, D. (1994), *Molecular Ecology of Rhizosphere Microorganisms*, pp. 67-89.

5. Furste, J.P., Pansegrau, W., Frank, R., Blocker, H., Scholz, P., Bagdasarian, M. and Lanka, E. (1986) Molecular cloning of the plasmid RP4 primase region in a multi-host-range tacP expression vector. *Gene*, **48**, 119-131.
6. Blumer, C., Heeb, S., Pessi, G. and Haas, D. (1999) Global GacA-steered control of cyanide and exoprotease production in Pseudomonas fluorescens involves specific ribosome binding sites. *Proc Natl Acad Sci U S A*, **96**, 14073-14078.
7. Ouahrani-Bettache, S., Porte, F., Teyssier, J., Liautard, J.P. and Kohler, S. (1999) pBBR1-GFP: a broad-host-range vector for prokaryotic promoter studies. *Biotechniques*, **26**, 620-622.
8. Ducret, V., Gonzalez, M.R., Leoni, S., Valentini, M. and Perron, K. (2020) The CzcCBA Efflux System Requires the CadA P-Type ATPase for Timely Expression Upon Zinc Excess in Pseudomonas aeruginosa. *Front Microbiol*, **11**, 911.
