## Supplementary material for "Global analysis of the zinc homeostasis network in *Pseudomonas aeruginosa* and its gene expression dynamics": Table S5

**Table S5:** Primers used in this study.

| Amplicon | PA number | Primer | Sequence 5'-3' | Position (from the AUG) | Length |
|---|---|---|---|---|---|
| **Coding regions** | | | | | |
| *czcC* | PA2522 | czcC1 | GGTCAGCATCGGCAGCAAGTACG | 834 | 206 |
| | | czcC2 | GGTCGTAGGCCTGTACCGCTTCG | 1039 | |
| *czcD* | PA0397 | 574 | GGCGTGGCCTTCTATATCCT | 285 | 183 |
| | | 575 | TCCAGACCTCCAGGTAGGC | 450 | |
| *ptrA / c* | PA2808 | 754 | CGTACCTTCACTGCCTCCAC | 502 | 176 |
| | | 755 | TTGTCCTTCTTCGTCGCTTC | 179 | |
| | PA2807 | 576 | GGTGCTCGGCGATATGTACT | 212 | 158 |
| | | 577 | GCATCTCCAGCATCTCCTTC | 350 | |
| *oprD* | PA0958 | oprD1 | ATCTACCGCACAAACGATGAAGG | 772 | 156 |
| | | oprD2 | GCCGAAGCCGATATAATCAAACG | 927 | |
| *oprF* | PA1777 | 594 | GGTTACTTCCTGACCGACGA | 172 | 209 |
| | | 595 | TCGCTGTTGATGTTGGTGAT | 380 | |
| *a* | PA2807 | 1338 | TTGACCTCGATCGCCCGAGG | 238 | 222 |
| | | 1339 | ACAGGATAGGCGACATCAAC | 35 | |
| *b* | PA2807-08 | 1340 | AGGCGGCGCGGAAACATACG | 70 | 318 |
| | | 1341 | ACCAGGAGAAACTGGCGAAG | 134 | |
| **Promoter regions** | | | | | |
| *pznuD* | PA0781 | 1254 | ACTTTCCCGTTCCTCTTCGC | -215 | 197 |
| | | 1255 | TGGGTCGGACTTTTTTCAGG | -18 | |
| *pPA1922* | PA1922 | 1256 | GAGCGCCATGATCTGCTCGG | -197 | 272 |
| | | 1257 | GAAGCAGGTAGAGGAACGGC | 75 | |
| *pPA2439* | PA2439 | 1344 | GTGCTGCACACAGACTCTCG | -140 | 167 |
| | | 1345 | GTCCAGGTCGACTCTCATCT | 17 | |
| *pPA2911* | PA2911 | 1258 | CATAGCAGGCCGGCGTGAAG | -458 | 229 |
| | | 1259 | TCACCGAAGTGACGCACCGG | -229 | |
| *pPA4063* | PA4063 | 1027 | GTTTCTCCACAAGCGAATCG | -4 | 182 |
| | | 1028 | ATGGGCCTGGGCGGCGAACG | -186 | |
| *pzrmA* | PA4837 | 1109 | GGCTGGGCTGGTCGTCGGA | -141 | 225 |
| | | 1110 | CCGCGGGACTTTCCGTTTCC | 84 | |
| *pznuA* | PA5498 | 969 | CTCTGCCAGACCAGTTCCAG | -227 | 214 |
| | | 970 | TGAAAGGTGGTTAGCGGGTA | -14 | |
| *prpmE2* | PA3600 | 1107 | TCGAGCAGCCGAATGGTCGC | -99 | 182 |
| | | 1108 | GGTGGAGCCGATCAGGAAGT | 83 | |
| *pdksA2* | PA5536 | 1105 | AGTAAAGGCGGAGGACGCGG | -137 | 203 |
| | | 1106 | AGAAGTCCTGCTGGGCTTCG | 66 | |
| **DNA cloning** | | | | | |
| **Feature** | | **Primer** | **Sequence 5'-3'** | **Length** | |

| | | | |
|---|---|---|---|
| *ΔPA2807* | 588 | GCCgaattcCATGTTCATCCCTCGTTACA | 489 |
| | 589b | GCCggtaccACATCTTTGTCAGCTTGCCG | |
| | 590b | GCCggtaccCGATCCGTATACACTTGCCG | 425 |
| | 591 | GCCggatccGGTCTTTCTGCAATACCGCG | |
| *ΔcopRS* | 715 | CCCgaattcCTGTCGGGAGCATTACATCT | 491 |
| | 716 | CCCggtaccTCGTTACATTTGCCGGCCAT | |
| | 717 | CCCggtaccAACGCGAAGCCCCCATGACG | 393 |
| | 718 | CCCggatccGTGCTGGTGGTGGTGCTTAC | |
| *pMMB66EH-PA2807* | 626 | GCCgaattcATGCTCCCGACAGCCAGCCG | 639 |
| | 627 | GCCggatccTTCACGCCGGCCGTCCGCCG | |
| *pME6001- PA2807::6His* | 1309 | CGCggatccTCAGCAGTTTCATGTTCATC | 1115 |
| | 1310 | CGCaagcttTCA**atgatgatgatgatgatg**GGGCTGCACGGTCAGTTGAC | |
| *pGEX2T-zur* | 1044 | GCGggatccATGTACAAGATTGCGCCCAAGACCC | 503 |
| | 1045 | GCCgaattcTCAGGCGTCCTTCTGGTCCC | |

Restriction sites are underlined (lowercase) and 6His tag is indicated as lowercase in bold
